# Comparison of long read sequencing technologies in resolving bacteria and fly genomes

**DOI:** 10.1101/2020.07.21.213975

**Authors:** Eric S. Tvedte, Mark Gasser, Benjamin C. Sparklin, Jane Michalski, Xuechu Zhao, Robin Bromley, Luke J. Tallon, Lisa Sadzewicz, David A. Rasko, Julie C. Dunning Hotopp

## Abstract

**Background:** The newest generation of DNA sequencing technology is highlighted by the ability to sequence reads hundreds of kilobases in length, and the increased availability of long read data has democratized the genome sequencing and assembly process. PacBio and Oxford Nanopore Technologies (ONT) have pioneered competitive long read platforms, with more recent work focused on improving sequencing throughput and per-base accuracy. Released in 2019, the PacBio Sequel II platform advertises substantial enhancements over previous PacBio systems.

**Results:** We used whole-genome sequencing data produced by two PacBio platforms (Sequel II and RS II) and two ONT protocols (Rapid Sequencing and Ligation Sequencing) to compare assemblies of the bacteria *Escherichia coli* and the fruit fly *Drosophila ananassae*. Sequel II assemblies had higher contiguity and consensus accuracy relative to other methods, even after accounting for differences in sequencing throughput. ONT RAPID libraries had the fewest chimeric reads in addition to superior quantification of *E. coli* plasmids versus ligation-based libraries. The quality of assemblies can be enhanced by adopting hybrid approaches using Illumina libraries for bacterial genome assemblies or combined ONT and Sequel II libraries for eukaryotic genome assemblies. Genome-wide DNA methylation could be detected using both technologies, however ONT libraries enabled the identification of a broader range of known *E. coli* methyltransferase recognition motifs in addition to undocumented *D. ananassae* motifs.

**Conclusions:** The ideal choice of long read technology may depend on several factors including the question or hypothesis under examination. No single technology outperformed others in all metrics examined.

## INTRODUCTION

Long read sequencing technologies enable the production of highly contiguous and accurate genome assemblies. Since the release of the Pacific Biosciences (PacBio) RS sequencer in 2011 and the Oxford Nanopore Technologies (ONT) MinION sequencer in 2014, improvements in sequencing chemistries and new sequencing platforms have continued to produce longer sequences and higher sequencing throughput, thus decreasing per-base sequencing costs [1]. Most recently, the PacBio Sequel II system advertises the highest throughput out of any of its sequencing platforms and includes two distinct sequencing modes: continuous long read sequencing (CLR) for ultralong reads and circular consensus sequencing (CCS/PacBio HiFi) for highly-accurate consensus reads. An insect genome assembled with Sequel II data has been published [2], however Sequel II comparative assembly performance relative to existing sequencing technologies is less clear.

Beyond improving genome assemblies, long read sequencing can be used as an alternative to bisulfite sequencing to detect genome-wide DNA methylation, producing the methylome of the organism. DNA methylation is found across the tree of life and is associated with a wide range of biological functions, including protection of host DNA against endonuclease cleavage, DNA replication, and gene expression [3]. DNA modification events are detected as measurements of DNA polymerase kinetics in PacBio SMRT sequencing [4–6] and as changes in the ionic current signal in the ONT nanopore [7, 8].

Although long reads can be useful in overcoming potential pitfalls of assembling with short read data alone, there are notable disadvantages of long read sequencing data. The error rate for single pass sequencing is ~13% for PacBio sequencers and ~15% for ONT [9, 10]. Methods have been developed to address these high error rates, such as the use of error correction before or after the assembly to achieve a highly accurate consensus sequence [11], or the derivation of a consensus read from multiple passes of a single template molecule during the sequencing run (*e.g.* PacBio HiFi; [12]). Artefactual chimeric reads, DNA sequences that originate from two distinct parent sequences, can also hinder the assembly process, although this is also a problem with all prior sequencing platforms. Chimeric reads have been reported in PacBio [13] and ONT sequencing [14], and preparations involving ligation and/or PCR amplification steps are likely to generate such artefacts.

Here, we investigate the quality of long read sequencing data produced using four methods: PacBio RS II, PacBio Sequel II CLR, ONT Rapid Sequencing Kit (ONT RAPID), and ONT Ligation Sequencing Kit (ONT LIG). We also sequenced Illumina and PacBio HiFi libraries for hybrid assemblies and genome polishing. To evaluate small genome assemblies, we used *Escherichia coli* E2348/69, a pathovar causing diarrheal illness with a complete genome sequence including numerous plasmids [15], making it an ideal reference for testing the completeness and accuracy of bacterial and plasmid assemblies. To compare assemblies of a larger genome, we produced long reads for *Drosophila ananassae* Hawaii which was previously sequenced but was highly fragmented [16, 17]. Overall, we demonstrate that no method was superior in all analyses performed, and the decision to use PacBio and ONT platforms for sequencing may depend on the specific question being addressed.

## RESULTS

### Read composition and *de novo* assembly performance in *E. coli*

*E. coli* sequencing data was generated from PacBio RS II, PacBio Sequel II, ONT RAPID, and ONT LIG (**Figure 1**; **Table S1)**. PacBio Sequel II libraries had the highest read N50 value of any library and represented a substantial improvement in read length and sequencing throughput relative to PacBio RS II. The ONT RAPID and LIG libraries had similar read N50 values (20-22 kbp) but 50-fold less total sequencing data was obtained from the RAPID run. The maximum read length for ONT libraries were about 50 kbp longer than PacBio Sequel II, representing a ~33% increase in these runs (**Table S1**).

**Figure 1.**
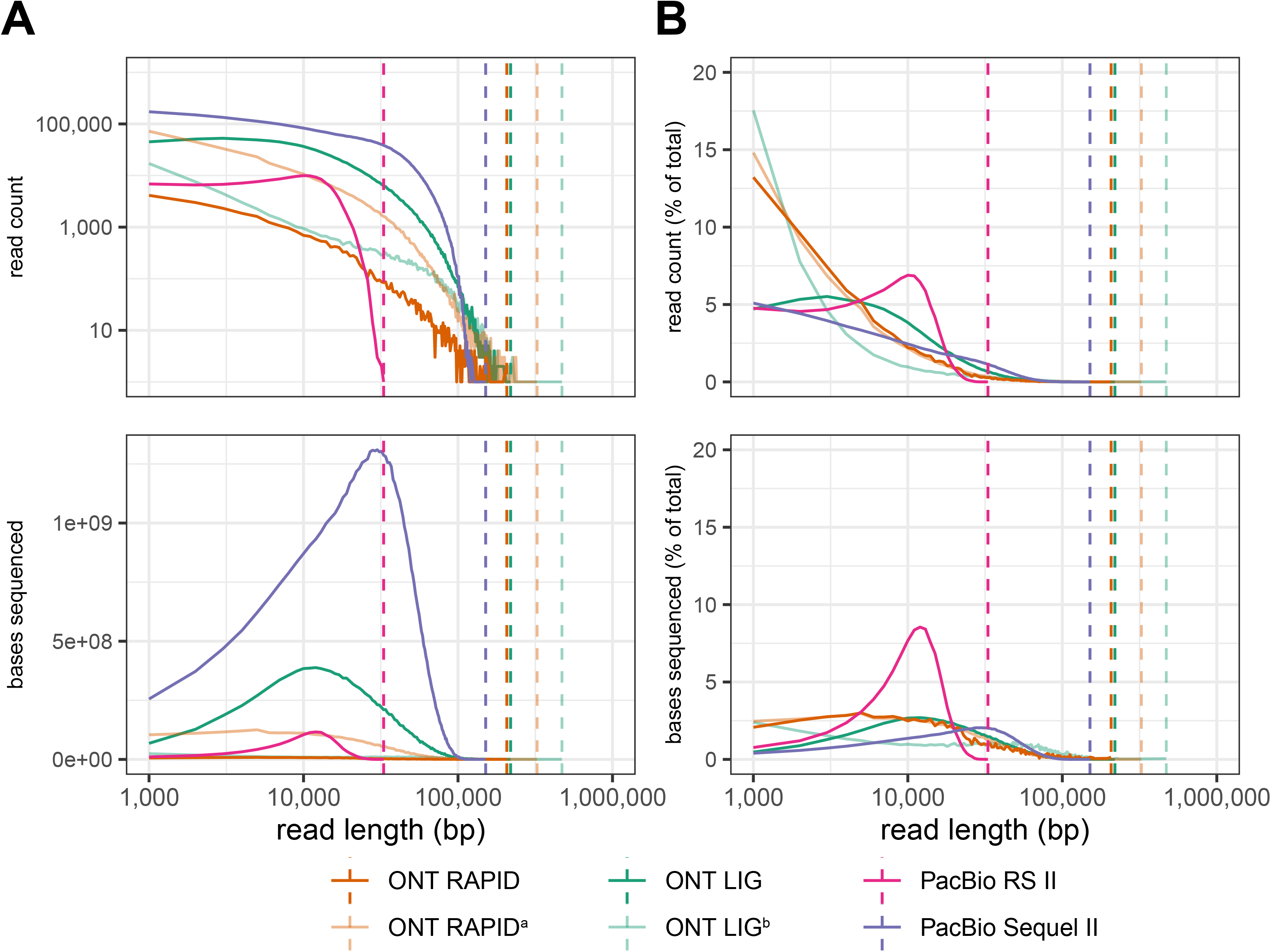
Read composition of *E. coli* long read libraries. Single end reads were placed into 1 kbp read length bins. Read counts and sequenced bases were calculated for each bin using the readlength.sh script of bbtools [50]. **A.** Absolute values for read counts and sequenced bases are plotted for each read length bin. **B.** Read counts and sequenced bases expressed as percentages of the total for each library are plotted for each read length bin. Vertical dotted lines correspond to maximum read length for each library. ^a^ONT RAPID sequencing run was sequenced following poor throughput of first RAPID run. ^b^ONT LIG sequencing run was performed without size-selection of DNA fragments.

Because we did not make an effort to sequence at the same depth in all libraries, the distributions of read lengths and sequenced bases per read length largely reflect differences in the total sequencing throughput of each run (**Figure 1A**). When read counts were expressed as percentages, the abundance of reads <5 kbp in ONT RAPID relative to other libraries reflects the lack of library size-selection in this protocol (**Figure 1B**). The PacBio RS II library had a large percentage of ~10 kbp reads with counts decreasing sharply to the library’s maximum read length of ~33 kbp. PacBio Sequel II had higher percentages at intermediate read lengths (20-100 kbp) but ONT had greater read representation at read lengths >100 kbp (**Figure 1B**). Additional ONT libraries were generated to assess variability in sequencing runs but were not assembled in this study.

To enable *de novo* assemblies using similar amounts of input sequencing data, read sets were randomly downsampled to approximate the PacBio RS II depth while maintaining the observed patterns for the full datasets (**Figure S1**). *E. coli* assemblies were produced from these random subsets (a) using Canu with long reads alone, (b) using Unicycler with long reads, and (c) using Unicycler with a hybrid approach combining the long reads combined with Illumina reads. All *E. coli* assemblies were ~5 Mbp, similar to the reference genome [15] but had variable length and contig composition in Canu and Unicycler long-read assemblies (**Table 1**; **Table S2**). Most libraries produced a single *E. coli* genome contig, whereas the ONT LIG Canu and Unicycler long read assemblies had two genome contigs, and the PacBio RS II Canu assembly had four (**Table 1**). All of the Unicycler hybrid assemblies using Illumina and long-read data produced identical 4,943,955 bp *E. coli* genome sequences and 5,218 bp p5217 plasmid sequences (**Table 1**; **Figure S2**). The 96,603 bp pMAR2 plasmid was identical in ONT RAPID, ONT LIG, and PacBio Sequel II hybrid assemblies, but different in the PacBio RS II hybrid assembly (96,184 bp). Contaminants were present in these assemblies that were removed following assembly; the Illumina libraries contain contaminating reads, as is common, and in this case included reads from human, *Neisseria gonorrhoeae*, and an unknown bacteria.

**Table 1.**
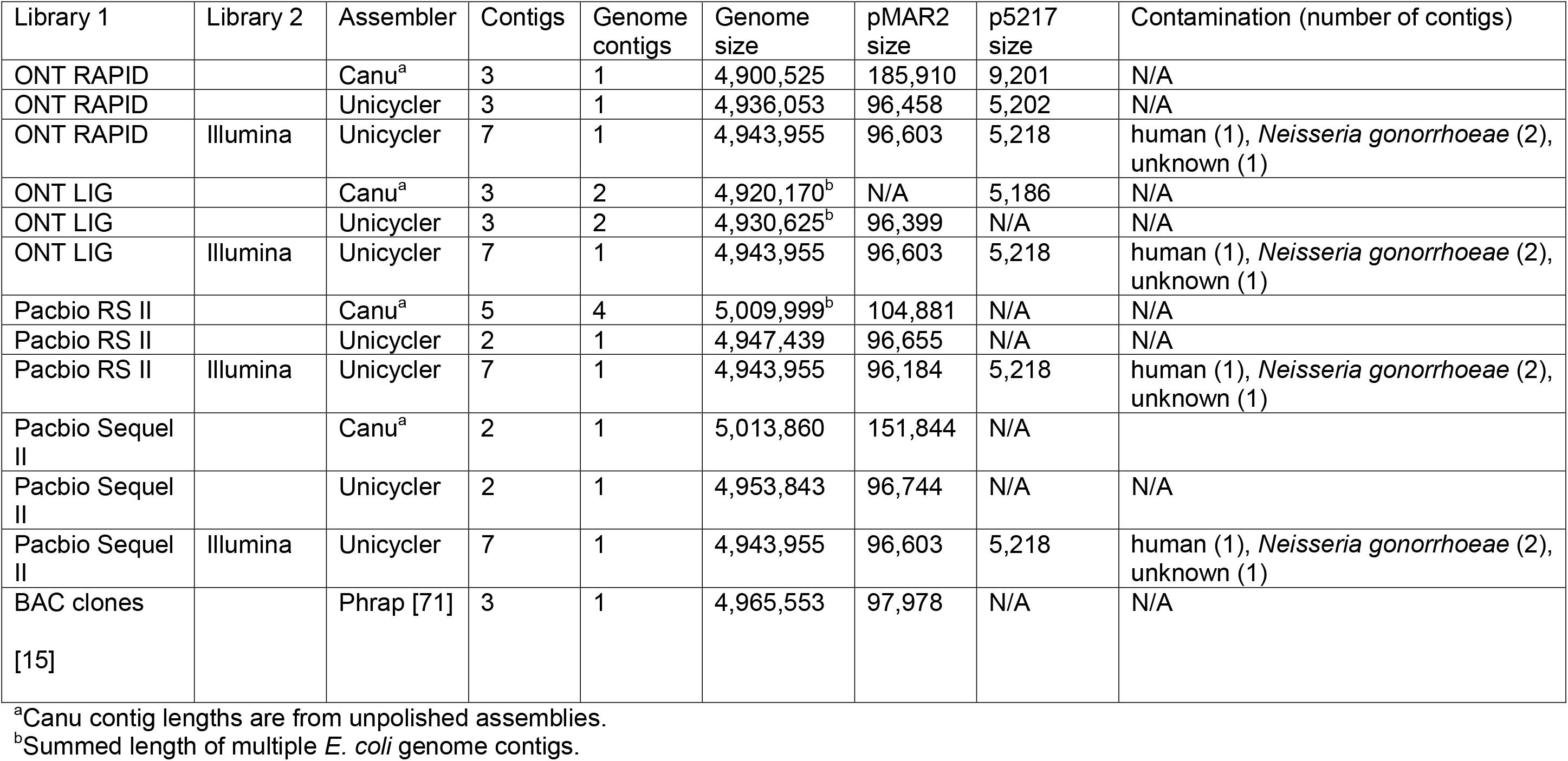
Summary of *E. coli* E2348/69 assemblies.

### Conserved gene content in *E. coli* assemblies

The presence of highly conserved bacterial genes in *E. coli* assemblies was assessed using BUSCO [18, 19]. In all cases, BUSCO failed to identify the 50S ribosomal protein L35 and 30S ribosome binding factor RbfA, which could be identified using BLASTN searches with protein queries from *E. coli* E2348/69 (Genbank CAS09394.1 and CAS10996.1, respectively). The remaining 146/148 BUSCO genes were characterized with variable success in long read assemblies. Many BUSCO genes were missing from assemblies of ONT reads without polishing, although Canu assemblies of only PacBio reads without polishing yielded all 146 genes. Polishing with Illumina or HiFi reads led to recovering all 146 genes in most cases, including: (a) all hybrid Unicycler assemblies that included Illumina sequencing data; (b) all ONT libraries with Illumina or PacBio HiFi reads; (c) PacBio RS II alone or with Illumina or PacBio HiFi reads; and (d) PacBio Sequel II data alone (**Table S2**). Sequel II assemblies with Illumina or PacBio HiFi reads were missing 2-15 additional genes, and many BUSCO genes were missing from the assemblies with Unicycler or Canu that used only long reads for polishing.

The consensus *E. coli* genome sequenced in this study was ~20 kbp shorter than the published genome sequence for this strain (**Table 2**). To validate the genome reduction, we focused on a ~16 kbp region that was present in the NCBI sequence but absent in the Unicycler assembly (**Figure S3A**). The junction spanning the deletion region was validated with PCR amplification (**Figure S3B**), and genes apparently absent from this *E. coli* specimen are involved in colanic acid biosynthesis (**Table S3**).

**Table 2.**
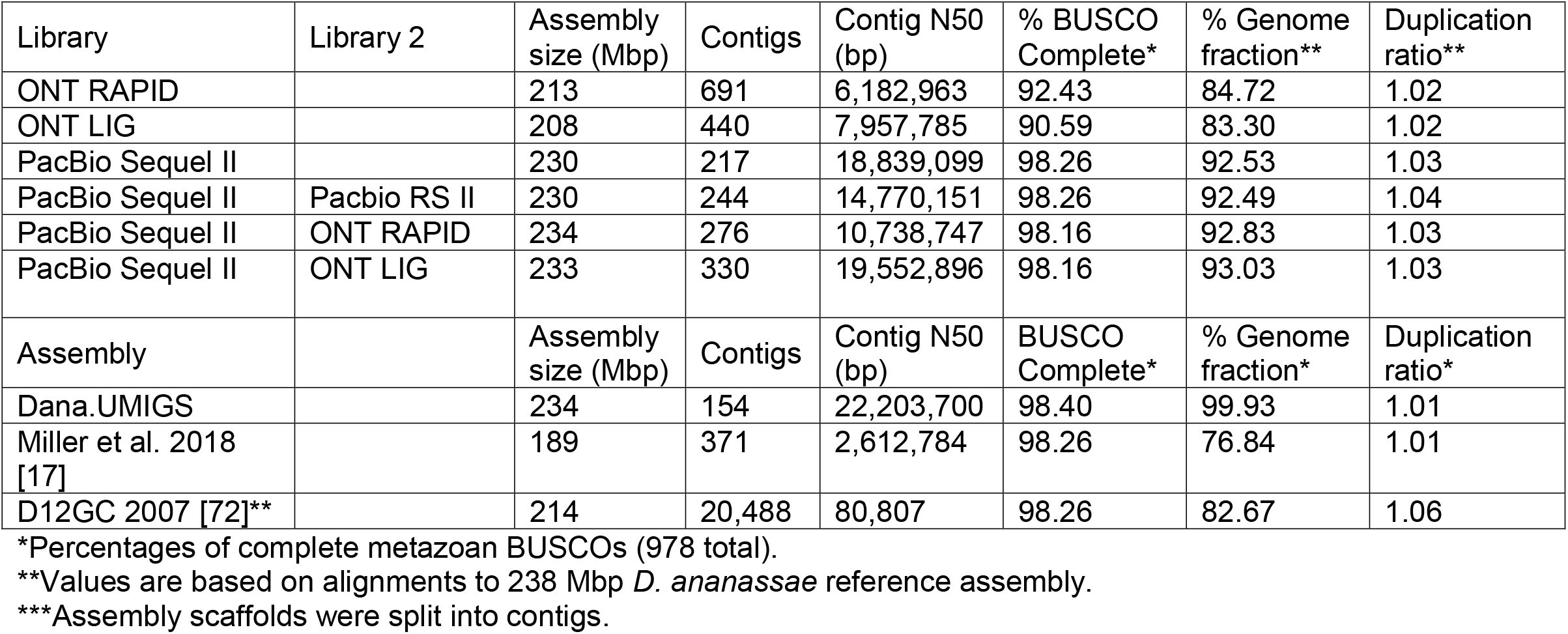
Summary of *D. ananassae* assemblies.

### Read and assembly correctness in *E. coli*

Assembly correctness was evaluated using a method similar to Wick et al. [20]. The *E. coli* chromosome was aligned to a trusted consensus that consisted of all ≥5 kbp contigs assembled separately using only Illumina data. Long read assemblies had 99-100% accuracy relative to the trusted consensus (**Table S2**). Assemblies polished with long read data alone were less accurate than those polished with short reads and PacBio HiFi reads. All Unicycler hybrid assemblies had 100% identity to the trusted consensus. ONT libraries assembled with Canu were the least accurate, and PacBio assemblies were consistently more accurate than ONT. Systematic, non-random errors in ONT sequencing data may cause this lower consensus accuracy [21].

The resolution of the *E. coli* genome into a single contiguous sequence enabled the assessment of the presence of chimeric reads in long read libraries. After mapping reads to the consensus Unicycler genome sequence, chimeras were quantified using Alvis [22]. The ONT RAPID library had the lowest percentage of putative chimeric reads (0.03%), while ONT LIG had the highest (1.44%), (**Table S4**). Visualization of alignments in IGV did not suggest an artificial inflation of chimeras (**Figure S4**).

### Detecting DNA modification using *E. coli* long read libraries

According to REBASE [23], *E. coli* E2348/69 has ten methyltransferases and three methylated motifs, including m6A modification of 5′-GATC-3′ by DNA adenine methyltransferase (Dam) and its three paralogs, m6A modification of 5′-ATGCAT-3 by YhdJ DNA methyltransferase, and 5mC modification of 5′-CCWGG-3′ by DNA cytosine methyltransferase (Dcm) [24].

Using the PacBio RS II library, the SMRT Tools DNA base modification pipeline identified five DNA motifs forming three distinct palindromes that were only enriched for m6A methylation (**Table S5**). There were 39,668 GATC sites identified in the *E. coli* dsDNA genome sequence, and >99.9% were characterized as methylated (**Table S5**), likely by Dam and its paralogs. Most GATC sites assigned as unmethylated were noncoding (**Table S6**), in agreement with the observation of methylase protection in noncoding regions in a previous study [25]. The YTCAN^6^GTNG/CNACN^6^TGAR motif had 878 sites with nearly ubiquitous methylation. This DNA methylase recognition sequence is shared with four REBASE entries, including three *E. coli* strains and *Shigella boydii* ATCC 49812 but was not previously characterized in this *E. coli* strain. The CYYAN^7^RTGA/TCAYN^7^TRRG motif had 579 sites and was nearly universally methylated but had no matches in REBASE (**Table S5**).

The greater sequencing depth of the Sequel II library resulted in higher modification quality values (QV) and GV/GT was a nondescript motif reported by default SMRT Tools parameters (**Table S5**). Guanine nucleotides had exceptionally high baseline modification QV, indicative of bias in the v0.9 beta sequencing chemistry likely resulting in spurious identification of GV/GT as methylated motifs (**Figure S5**). This bias was not unexpected since base modification was not supported in this chemistry release during the beta testing phase when this sequencing was completed. However, at the suggestions of Pacific Biosciences, increasing the modification QV threshold removed the GV/GT motifs from the report but identified fewer methylated GATC sites (99.3%; **Table S5**). Elevated baseline modification QV scores were not observed in RS II data (**Figure S5**).

DNA methylation in *E. coli* ONT libraries was assessed with tombo [26]. Tombo uses canonical base models for *de novo* detection of DNA methylation events in addition to alternative models that are modification-centric (*e.g.* m5C, m6A) or motif-centric (*e.g.* GATC, CCWGG). The *de novo* model identified a high incidence of methylation activity at the GATC, YTCAN^6^GTNG, and TCAYN^7^TRRG palindromic motifs similar to the PacBio sequencing (**Figure 2A; Figure S6**).

**Figure 2.**
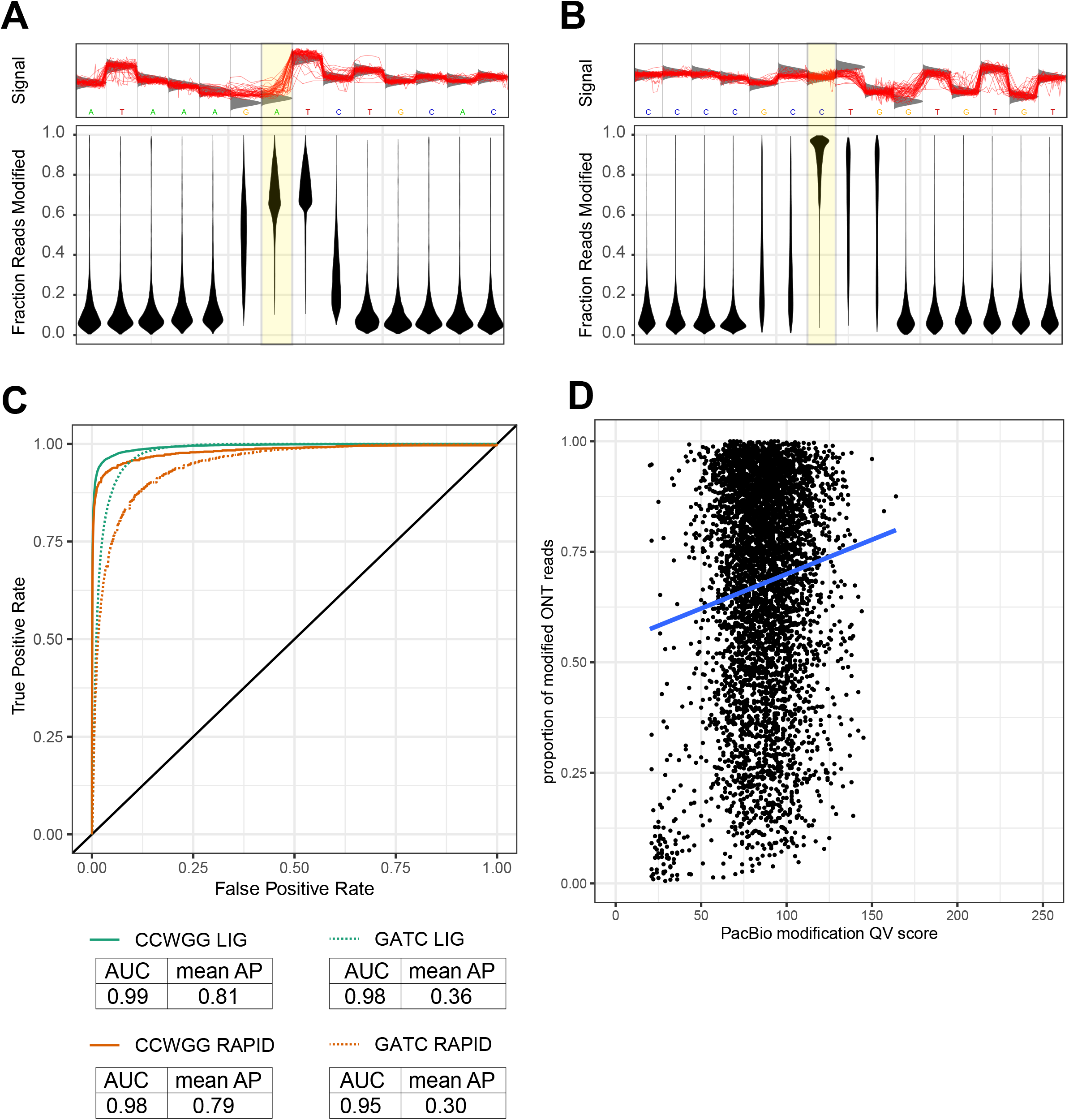
Evidence of DNA methylation in *E. coli* E2348/69 using long read sequencing. Methylation at A. GATC and B. CCWGG motifs are supported using ONT LIG sequencing. Top: An example motif is shown, with individual reads plotted to the region shown in red. The expected raw signal distribution using a canonical base model (=unmethylated DNA) is shown in grey. The location of known methylation in *E. coli* is highlighted. Bottom: The fraction of reads supporting a modification event is reported for each position in the motif, and the distribution of proportions are shown. Higher values indicate the motif is more ubiquitously methylated in the *coli* genome. Distributions are shown for 11,313 GATC motifs and 20,063 CCWGG motifs. C. ROC curves for detection of methylation at known motifs. GATC and CCWGG motifs were considered ground truth and modified base statistics of these sites were compared against statistics at other base modification sites. ROC curves for ONT RAPID (~60X depth) and ONT LIG (~3,280X) are plotted with AUC and AP values for each condition shown. D. Association of m6A modifications assessed using PacBio and ONT sequencing. All adenine sites meeting the minimum modification QV reporting threshold of 20 were cross-referenced for corresponding dampened fraction values in ONT LIG sequencing. A random sample of 5,000 adenine sites are plotted (total = 43,783). A linear regression was fitted to the data.

There was no apparent support for methylation at ATGCAT motifs, which is consistent with the PacBio results (**Figure S6**). Unlike PacBio, ONT sequencing identified cytosine methylation at CCWGG motifs (**Figure 2B**). Detection of methylated GATC and CCWGG motifs improved with the increased sequencing depth of the ONT LIG data, supporting previous findings (**Figure 2C**) [26].

Due to the distinct pipelines of SMRT Tools and tombo, it is difficult to assess the convergence of DNA modification using PacBio and MinION sequencing. In addition to the joint identification of specific m6A-methylated motifs, there is a positive association between PacBio modification QV and the proportion of ONT reads supporting a methylation event when considering all *E. coli* adenine residues (**Figure 2D**).

### *E. coli* plasmids are underrepresented in ligation-based libraries

The presence of the ~97 kbp plasmid pMAR2 and the ~5 kbp plasmid p5217 was confirmed with searches against the plasmid database PLSDB [27]. Without polishing, the plasmid contig was frequently up to 2X-longer than the actual plasmid, but this could be resolved with polishing and circularization (**Table S2**). The p5217 plasmid was not assembled with only PacBio RS II or Sequel II data, the pMAR2 was not assembled with ONT LIG data alone, and the previously reported pE2348-2 plasmid from this *E. coli* strain (Genbank FM18070.1) [15] was not present in any assemblies.

Size-selected libraries may fail to produce plasmid sequences in assemblies if plasmid sizes are small, but assemblers could also fail to properly identify plasmids because of k-mer abundance differences due to copy-number differences. To differentiate between these scenarios, reads were mapped to a reference containing the consensus *E. coli* genome and two plasmids from the Unicycler assemblies with Illumina error correction that was recovered independently four times, once for each long read library. The combined depth of pMAR2 and p5217 contributed to nearly 5% of total sequencing depth in ONT RAPID libraries with both being estimated to be present in 2-3X copy number relative to the *E. coli* genome (**Table S7**). Sequence reads from plasmids were less abundant in other libraries, contributing to less than 2% of total sequencing depth in ONT LIG and PacBio datasets. The size-selected PacBio libraries had no primary alignments reads mapping to the p5217 plasmid.

Using qPCR, there are approximate two copies of p5217 per genome, similar to the observed ratio in the ONT RAPID reads (**Table S7**; **Table S8**; **Table S9**; **Figure S7**) while there is approximate one copy of pMAR2 per genome. Assuming that the qPCR results are correct, this suggests that it is overrepresented in the ONT RAPID library and underrepresented in ONT LIG and PacBio libraries, although amplification biases have been demonstrated in plasmid qPCR experiments [28]. The results for the pE2348-2 replicates resembled the negative control, suggesting this plasmid is not present in this *E. coli* sample (**Figure S7**). These findings are consistent with previous observations of underrepresentation of small plasmid sequences in size-selected libraries [20, 29].

### ONT variability

There are reports of ONT sequencing variability, which was assessed with an additional ONT RAPID library as well as an additional ONT LIG library that lacked shearing and size selection steps. The second ONT RAPID run generated 7.2 Gbp of sequencing data (an 8-fold increase after correcting for run time) and had a similar read length distribution as the first run (**Table S1**; **Figure 1**). The second ONT LIG run generated ~1.0 Gbp of sequencing data and had a larger proportion of short reads sequenced, similar to the two ONT RAPID runs (**Table S1**; **Figure 1**). Estimated copy number of both plasmids was similar in the two ONT RAPID runs (**Table S7**). Recovery of plasmid reads was superior in the second ONT LIG library, with pMAR2 copy number closer to the expected 1X but p5217 remaining underrepresented (**Table S7**). Overall, all but ONT RAPID gave lower than expected plasmid sequencing depth ratios while ONT RAPID was the only protocol to consistently produce reads corresponding to small plasmids.

### Read composition and *de novo* assembly performance in *D. ananassae*

*D. ananassae* sequencing data produced from PacBio and ONT libraries had read composition profiles similar to *E. coli* (**Table S1**; **Figure S8**). The PacBio Sequel II library had the highest read N50 value while ONT libraries had longer maximum read lengths. The two ONT libraries had similar overall throughput, with RAPID having higher representation of <10 kbp reads and LIG having higher representation of >10 kbp reads (**Figure S8**). The sequencing depth of the PacBio RS II library (~4X) was too low to assemble individually.

*D. ananassae* assemblies were produced (a) using Canu with individual long read libraries and (b) using Canu to generate hybrid assemblies with combined read data from PacBio Sequel II and one additional read library. All assemblies were generated with default Canu parameters, with the longest reads up to ~40X sequencing depth used to construct the assembly. *D. ananassae* assembly sizes were 208-234 Mbp, partially overlapping the 185-224 Mbp range of reported genome size estimates for this species (**Table 2**) [30]. All assemblies had fewer than 700 contigs and had maximum contig sizes >20 Mbp. Among single library assemblies, the PacBio Sequel II assembly had the lowest contig count and the highest contig N50. Despite ONT reads being the longest, the hybrid ONT RAPID + Sequel II assembly had a lower N50 value and the largest contig was smaller (**Figure S9; Table S10**). The hybrid Sequel II + RS II had the largest single contig of any assembly, however other large contigs corresponding to euchromatic arms were smaller in this assembly relative to Sequel II alone and Sequel II + ONT LIG (**Figure 4**). All assemblies produced in this study were more contiguous than prior assemblies using capillary-based sequencing [16] and ONT sequencing [17] (**Figure S9; Table S10**).

### Comparison of major chromosomes in *D. ananassae* genome assemblies

The *D. ananassae* genome contains three euchromatic chromosomes (X, 2, 3) and two heterochromatic chromosomes (4, Y) [31]. Since all assemblies were more contiguous than previously published *D. ananassae* genomes, a new high-quality *D. ananassae* reference genome was separately assembled (referred to here as Dana.UMIGS) to facilitate comparisons between assemblies generated in this study. Briefly, the Dana.UMIGS assembly was constructed by merging a Flye assembly generated using Sequel II reads with a Canu assembly using Sequel II + ONT LIG reads and subsequently removing contaminants, assembly artefacts, and erroneously duplicated regions. The final Dana.UMIGS genome assembly was 234 Mbp, had 154 contigs, and resolved the six euchromatic arms (XL, XR, 2L, 2R, 3L, 3R) into six contigs totaling 158 Mbp (**Figure 3–Figure 5**). The spatial organization of the euchromatic arms is supported by physical maps of polytene chromosomes, including a known strain-specific chromosomal inversion on 3L (**Figure 3**–**Figure 5; Table S11**) [31]. Chromosome arms were more highly fragmented in assemblies using individual long read libraries, with these same regions in at least 11, 15, and 26 contigs in the PacBio Sequel II, ONT LIG, and ONT RAPID assemblies, respectively (**Figure 3**–**Figure 5; Table S12**–**Table S14**). Chromosome X was more fragmented in ONT assemblies and the order and orientation of the X pericentromere regions were not consistently resolved across assemblies (**Figure 3**). The euchromatic arms of chromosomes 2 and 3 were more contiguous and the Sequel II assembly contained pericentromeric regions in these chromosomes (**Figure 4**; **Figure 5**). Differences in chromosome fragmentation in individual library assemblies enabled the resolution of contiguous sequence in the Dana.UMIGS assembly which used both ONT and PacBio reads. Chromosomes 2L and 2R were resolved from multiple contigs in the ONT LIG and PacBio Sequel II assembly, respectively (**Figure 4**) and the entire chromosome 3 was assembled into a single large contig (**Figure 5**).

**Figure 3.**
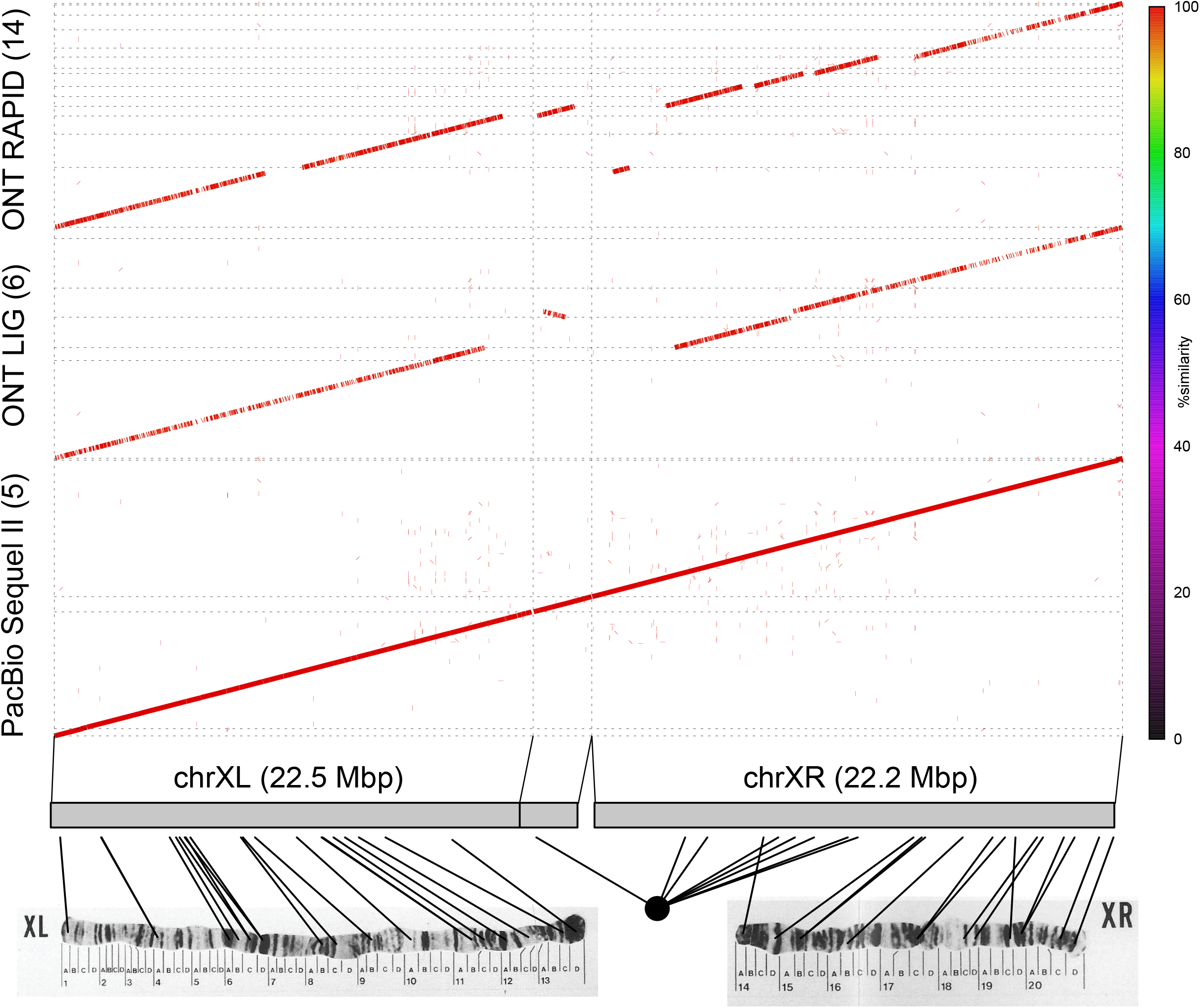
Alignments of contigs from long read assemblies to polytene maps of *D. ananassae* chromosome XL and XR. Top: Alignments of Dana.UMIGS chromosome X contigs to corresponding contigs generated from individual long read sequencing libraries. Dot plots were generated using nucmer in the MUMmer3 package. Parentheses on the y-axis indicate the number of identified contigs contributing to chromosome arms. Bottom: contigs from the Dana.UMIGS assembly are labeled, and lines connect polytene map coordinates with estimated locus positions generated with BLAST searches. Original images for polytene maps are from [73]. Permissions for the use of polytene map images were purchased from Karger Publishers.

**Figure 4.**
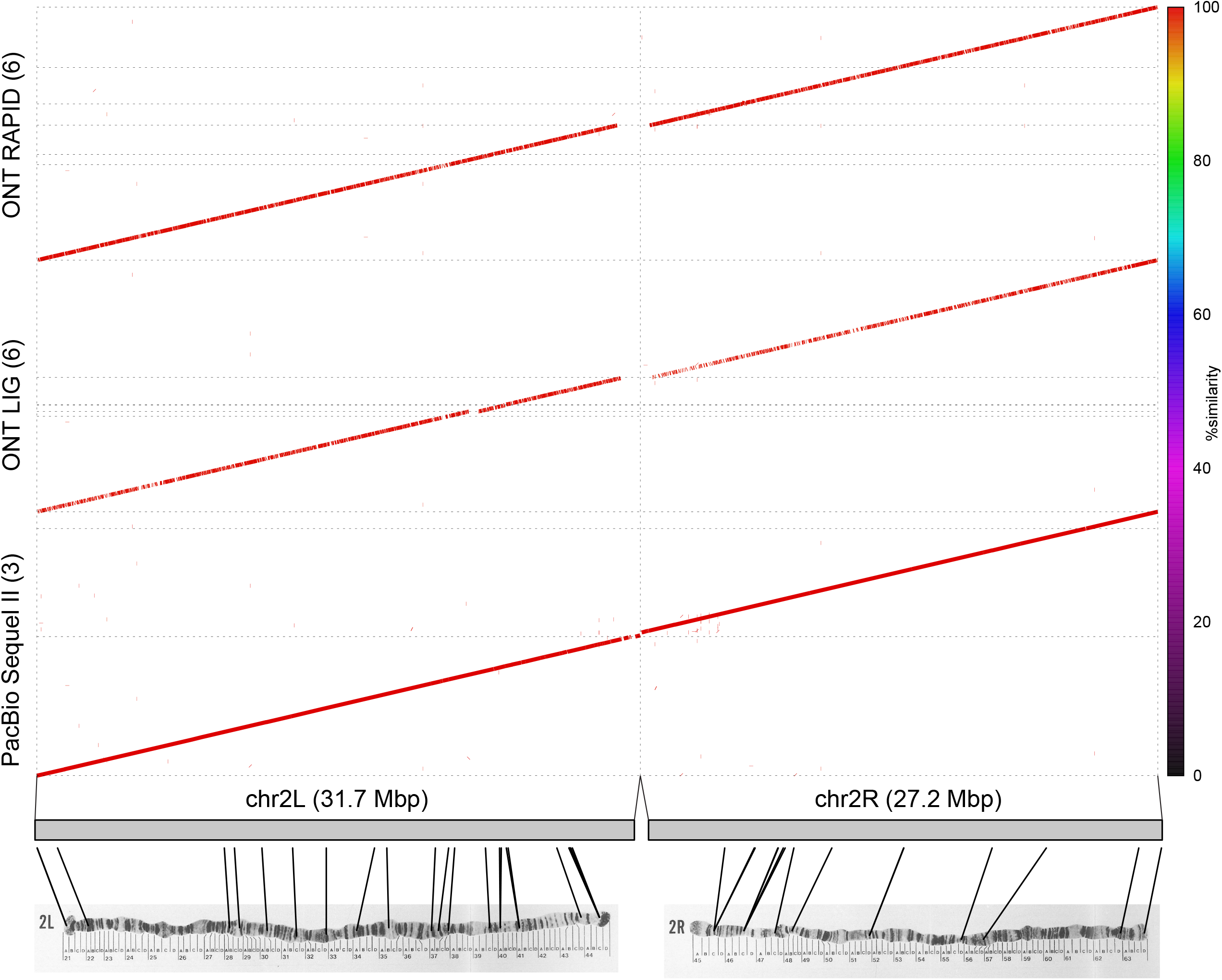
Alignments of contigs from long read assemblies to polytene maps of *D. ananassae* chromosome 2L and 2R. Top: Alignments of Dana.UMIGS chromosome 2 contigs to corresponding contigs generated from individual long read sequencing libraries. Dot plots were generated using nucmer in the MUMmer3 package. Parentheses on the y-axis indicate the number of identified contigs contributing to chromosome arms. Bottom: contigs from the Dana.UMIGS assembly are labeled, and lines connect polytene map coordinates with estimated locus positions generated with BLAST searches. Original images for polytene maps are from [73]. Permissions for the use of polytene map images were purchased from Karger Publishers.

**Figure 5.**
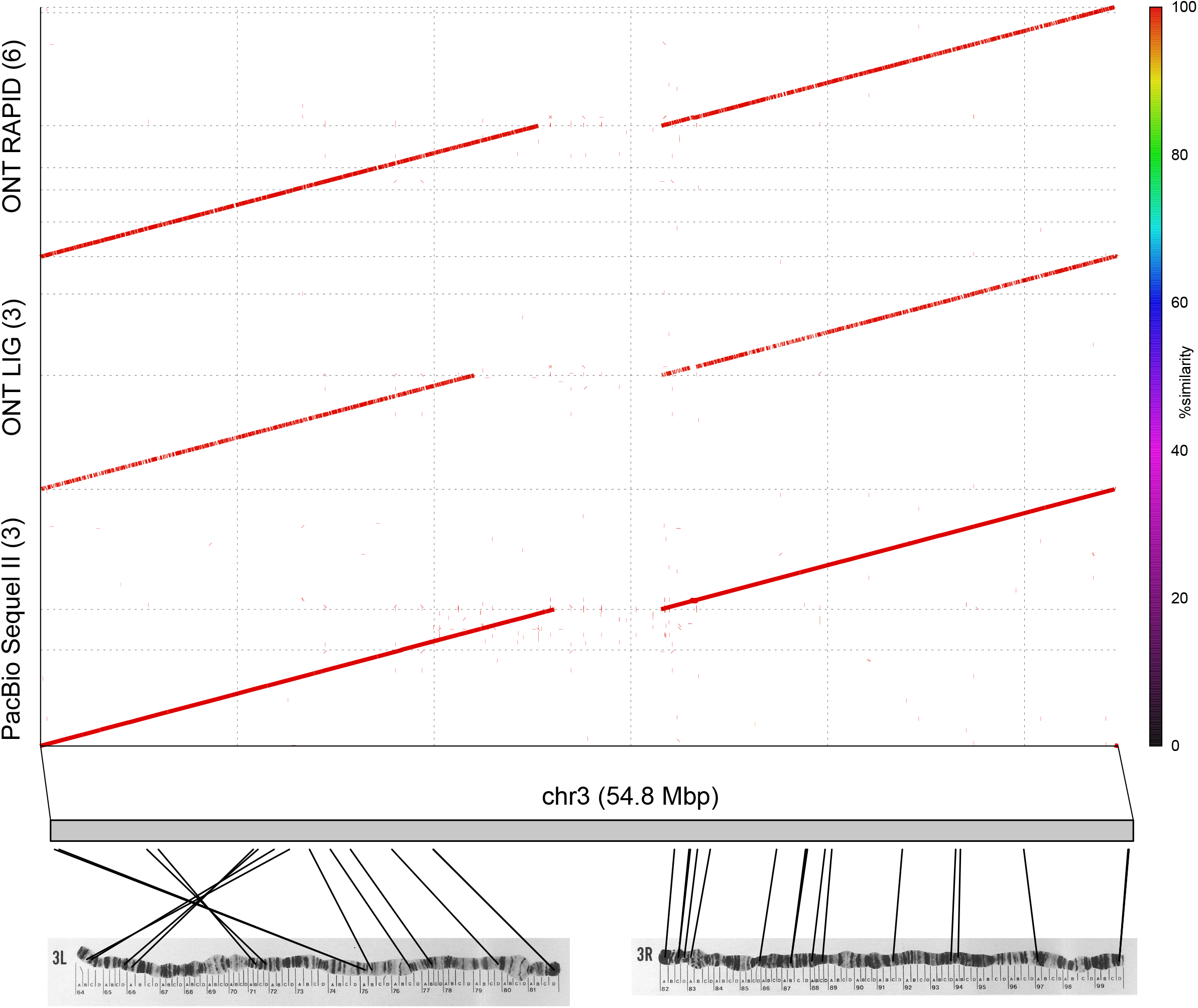
Alignments of contigs from long read assemblies to polytene maps of *D. ananassae* chromosome 3L and 3R. Top: Alignments of the Dana.UMIGS chromosome 3 contig to corresponding contigs generated from individual long read sequencing libraries. Dot plots were generated using nucmer in the MUMmer3 package. Parentheses on the y-axis indicate the number of identified contigs contributing to chromosome arms. Bottom: contigs from the Dana.UMIGS assembly are labeled, and lines connect polytene map coordinates with estimated locus positions generated with BLAST searches. Original images for polytene maps are from [73]. Permissions for the use of polytene map images were purchased from Karger Publishers.

### Conserved gene content in *D. ananassae* assemblies

The identification of complete metazoan BUSCOs was consistently high (90-98%) across *D. ananassae* assemblies (**Table 2**; **Table S10**). As observed in *E. coli*, ONT assemblies had a higher proportion of missing and fragmented BUSCOs while PacBio assemblies had more successfully characterized single-copy BUSCOs (**Table S10**). After polishing the ONT LIG + PacBio Sequel II assembly, maximal BUSCO recovery (962/978; ~98%) was produced with PacBio HiFi polishing, similar to the proportion found in the Dana.UMIGS assembly (963/978) and the Miller *et al*. genome assembly (961/978) (**Table S10**). Of the 14 genes characterized as incomplete by BUSCO, 13 were incomplete in the published Miller *et al*. assembly and eight were incomplete in *D. melanogaster* Release 6 (Genbank GCA_000001215.4), meaning that these genes may indeed be absent from *D. ananassae*.

### Assembly correctness in *D. ananassae*

Assemblies using PacBio Sequel II data were ~10-20 Mbp larger relative to ONT assemblies (**Table 2**). The genome fraction was calculated for each assembly in QUAST-LG as the length of aligned sequence in a queried assembly divided by the Dana.UMIGS assembly size. The genome fraction was lower in ONT versus PacBio assemblies (**Table 2**). The duplication ratio was calculated for each assembly in QUAST-LG as the length of aligned sequence in a queried assembly divided by the length of aligned sequence in Dana.UMIGS. PacBio assemblies had slightly higher duplication ratios relative to ONT assemblies (**Table 2**). The number of duplicated BUSCOs was also 3-4X higher in PacBio assemblies compared to ONT assemblies and published *D. ananassae* genomes (**Table S10**). While there may be true duplications in *D. ananassae*, the higher duplicated BUSCOs could be a consequence of mis-assembled regions due to elevated chimeric reads as observed in *E. coli*. The sequenced *D. ananassae* line was highly inbred, therefore duplicated regions are less likely due to recent divergence of haplotypes and more likely due to sequencing/assembly errors.

The Dana.UMIGS assembly was polished for successive rounds with PacBio Sequel II and PacBio HiFi data and was compared to other assembled genomes to evaluate assembly accuracy using QUAST-LG. Consensus accuracy was estimated by dividing 100 kbp by the sum of the mismatch rate (MMR) and the indel rate (IDR), both which were calculated in QUAST-LG per 100 kbp of aligned sequence. All assemblies had >99% consensus accuracy (**Table S10**). PacBio assemblies had fewer mismatches (29-90 MMR) than ONT (134 and 167 MMR in ONT LIG and RAPID, respectively) and indel frequencies in PacBio assemblies were tenfold lower (**Table S10**). Although we did not exhaustively test genome polishing schemes for all *D. ananassae* assemblies, polishing the ONT LIG + PacBio Sequel II using various read sets reduced the frequency of mismatches and indels relative to the original assembly, and the highest consensus accuracy (>99.9%) was achieved with PacBio HiFi polishing.

### Assembly of heterochromatic regions in *D. ananassae*

Repeat-rich regions are notoriously difficult to assemble and thus tend to be understated in genome reports. Substantial portions of the *D. ananassae* genome are highly heterochromatic, including all of chromosomes 4 and Y. *D. ananassae* has an expanded chromosome 4 relative to other *Drosophila* species, appearing similar in size to the X chromosome [31]. The increased size of chromosome 4 is partially attributed to its incorporation of large lateral gene transfers (LGT) from its *Wolbachia* endosymbiont (*w*Ana) [32, 33]. There is also evidence for retrotransposon proliferation in *D. ananassae* chromosome 4 relative to *D. melanogaster* [34].

The sequencing of long reads spanning repeat regions should enable the resolution of heterochromatin in fewer and longer contigs. To compare the contiguity of heterochromatic regions between assemblies, an additional QUAST-LG analysis was performed after removal of sequences corresponding to the euchromatic chromosome arms. The 206 heterochromatic contigs in the PacBio Sequel II assembly had a size of 76 Mbp and a contig N50 of 1.6 Mbp (**Table S15**). ONT assemblies had over twice the number of heterochromatic contigs and N50 values <750 kbp, consistent with the lower overall contiguity in ONT assemblies (**Table S15**).

### Detecting DNA modification using *D. ananassae* long read libraries

The extent of DNA methylation in *Drosophila* is not fully understood. Methylation in animals consists primarily of cytosine modification in CpG islands maintained by multiple DNA methyltransferases (DNMTs) [35]. *D. melanogaster* lacks homologs of DNMT1 and DNMT3 but possesses DMNT2 which is highly conserved across dipterans, mouse, and human [36]. In the attempt to characterize methylation in *D. melanogaster*, multiple studies disagree in (a) the extent of genome-wide cytosine methylation and (b) the role of DMNT2 in this process [37–40]. The retention of cytosine methlylation in DNMT2-knockout embryos [41, 42] indicate there are unidentified methyltransferases in *Drosophila*. Global 5mC has been quantified in *D. ananassae* using liquid-chromatography-mass spectrometry [43], but methylated motifs have not been reported.

Given its lower sequencing depth, the application of DNA modification pipelines is less reliable in *D. ananassae*. The PacBio IPD signatures of 5mC modifications are more challenging to detect relative to 6mA and require >250X depth or enzymatic conversion of 5mC to improve detection [44, 45]. Therefore, not surprisingly, we could not identify methylation signatures in *D. ananassae* using PacBio libraries. The <40X depth in ONT libraries was too low for robust genome-wide methylation calls. The Tombo 5mC model-based calling of the ONT LIG library permitted preliminary analysis: The 1000 regions in the ONT LIG + Sequel II assembly with the highest proportion of ONT reads supporting DNA methylation contained CG and GC dinucleotides (**Data S1**), however the precise methylation sites cannot be readily identified in more complex motifs using this method.

## DISCUSSION

Our study demonstrates that highly contiguous assemblies can be obtained with long-read technologies, but highly accurate assemblies benefit from error correction with more accurate reads than currently available long reads alone. All *E. coli* assemblies surpassed 99% accuracy when using long read data alone, and accuracy was further improved when using Illumina or PacBio HiFi data for hybrid assembly and error correction. *D. ananassae* assemblies had improved contiguity relative to published genomes, and nearly all euchromatic chromosome arms were resolved in single contigs.

### Comparative analyses support superior performance of PacBio Sequel II libraries

PacBio Sequel II sequencing represents a major advancement in sequencing throughput over previous PacBio platforms with the production of more sequencing data and longer reads versus RS II and the Sequel I (not tested here). Although ONT libraries had longer reads sequenced, Sequel II had a larger pool of ultra-long reads, demonstrated by higher read N50 values. Additionally, the overall sequencing throughput of ONT was more variable: the runs produced between 12-670 Mbp/hr sequencing data, compared to 224-340 Mbp/hr in PacBio RS II and 2.1-2.7 Gbp/hr in PacBio Sequel II.

While greater sequencing depth increases the likelihood of producing high consensus accuracy resolution from error-prone reads, the subsampling of datasets confirmed that *E. coli* assemblies using PacBio sequencing were the most accurate. Highly conserved bacterial genes were also more consistently characterized in PacBio assemblies. Consensus accuracy and BUSCO identification were not improved when using long reads alone for polishing, suggesting that the sequencing depth of corrected reads was too low and/or that persistent errors remain in long reads following read correction with Canu. Superior accuracy was achieved after polishing with Illumina or PacBio HiFi reads for polishing, although residual errors in repetitive regions may remain when using these datasets for correction that won’t be assessed with this BUSCO-based method.

*D. ananassae* assemblies using PacBio Sequel II data were the most contiguous despite the sequencing of longer ONT reads. Heterochromatic regions were also more contiguous in PacBio assemblies. Since assemblies were generated using default Canu parameters, *i.e.* assembling the longest reads up to ~40X sequencing depth, differences in assembly contiguity could be partially attributable to the increased percentage of bases sequenced in ultra-long reads in Sequel II. However, higher incidence of errors in ONT reads could also hinder overlap detection. Although the summary statistics commonly used to describe genome assemblies (*e.g.* contig count, contig N50, maximum contig length) were superior in PacBio, these assemblies might also have more duplicated content from uncollapsed regions; researchers must exercise appropriate caution when choosing the “best” assembly to report.

### PacBio sequencing is not optimal for all sequencing applications

While PacBio Sequel II demonstrated the best results for most data quality tests, there are certain situations where alternatives to PacBio sequencing might be preferred. First, the upper limit of PacBio read length is determined by polymerase processivity and DNA loading into the SMRT Cell, meaning ONT sequencing can span longer repetitive regions. In highly repetitive eukaryotic genomes a hybrid approach might be warranted; multiple *D. ananassae* assembly gaps were closed using both PacBio and ONT. Second, size-selected libraries will exclude small sequences from the assembly. Our *E. coli* results demonstrated the complete omission of a 5 kbp plasmid and the underrepresentation of a 97 kbp plasmid using Sequel II sequencing. Conversely, ONT RAPID produced plasmid sequencing data much closer to expected proportions. Third, the higher cost of entry for the PacBio platform might be prohibitive for some researchers, despite the lower cost per base for Sequel II sequencing. Fourth, chimeric reads are more common in library preparations that involve ligation. Unicycler uses long reads to generate scaffold bridges across contigs assembled with Illumina data, meaning assemblies shouldn’t be negatively impacted by chimeric reads [20, 46]. Although the overall frequency of chimeric reads is low, additional investigation of the occurrence and genome-wide distribution of chimeras is needed, particularly for eukaryotic genomes. Fifth, characterization of cytosine methylation is more challenging with PacBio sequencing.

### Development of new sequencing products and bioinformatics tools will continue to improve long read sequencing

The rapid turnover of sequencing platforms and analysis pipelines will continue to improve the utility of long read sequencing data. Since the design of this experiment, Oxford Nanopore Technologies have begun distribution of the R10 nanopore which advertises to improve consensus accuracy to over 99.99%. New Sequel II sequencing chemistry released by PacBio claims to improve performance, including the reduction of baseline DNA modification scores. The increased use of long read sequencing data has spurred a plethora of bioinformatic tools for long read overlap detection, contig assembly, and error correction [11, 47, 48]. Platform-specific tools have been developed to achieve optimal results given the underlying features of long read data (*e.g.* Arrow and Nanopolish used for polishing genomes using PacBio and ONT data, respectively). After generating assemblies for this study, Canu v.1.9 has been released allowing users to set parameters specific to PacBio HiFi sequencing. While it is possible that improved results could have been obtained in this study by using platform-specific tools, we chose tools for the study on the basis of a) their wide usage in long read genome assembly and b) their platform independence. Nevertheless, it is possible that the tools with the specified parameters are better able to handle the error profile of data from a specific platform, leading to the observation of superior performance in many of our tests.

## CONCLUSIONS

With the arrival of PacBio Sequel II, researchers can achieve unprecedented throughput in long read sequencing data. The advancement of Sequel II confers an increase in consensus accuracy and a higher likelihood of sequencing across repetitive regions. Increased adoption of long read sequencing platforms promises to revolutionize genomics research.

## METHODS

### Biological samples

*Escherichia coli* E2348/69 cultures grown overnight in L-broth were pelleted (12,000 x g), resuspended in 50 mM Tris, 1 mM EDTA, 10 μl RNAse (20mg/ml) and lysed with 0.4% SDS (final) at 56 °C for 30 min. A 0.5 volume of 7.5 M ammonium acetate was added, and samples were incubated for 15 min on ice. Genomic DNA was extracted with phenol:chloroform:isoamyl alcohol followed by chloroform: isoamyl alcohol and precipitated with isopropanol. After two washes with 70% ethanol, the pellet was allowed to air dry and resuspended in water.

*Drosophila ananassae* Hawaii (14024–0371.13) were obtained from the *Drosophila* Species Stock Center (University of California, San Diego, USA). Populations were grown on Jazz-Mix *Drosophila* food (Applied Scientific) in plastic bottles at 25°C and 70% humidity with a 12hr-12hr light-dark cycle. Flies were treated with tetracycline to remove the *Wolbachia* endosymbiont. Genomic DNA was extracted from ~350 flies with phenol:chloroform:isoamyl alcohol followed by chloroform: isoamyl alcohol and precipitated with isopropanol. After two washes with 70% ethanol, the pellet was allowed to air dry and resuspended in water.

Genomic DNA was quantified using the Qubit 4 fluorometer (Thermo Fisher Scientific) and the presence of >20 kbp fragments were validated using the FEMTO Pulse automated pulsed-field capillary electrophoresis instrument (Agilent Technologies).

### Nanopore libraries and sequencing

ONT RAPID libraries for *E. coli* and *D. ananassae* were prepared with the Rapid Sequencing Kit SQK-RAD004 (Oxford Nanopore Technologies) using 10 μL DNA, 8.5 μl EB, 1.5 μl FRA, and omitting library-loading beads. After adding Rapid adapters, the reactions were incubated for 30 min at room temperature. One 24 hr sequencing run was performed for *E. coli* and two sequencing runs were performed for *D. ananassae* using FLO-MIN106 R9 MinION flowcells (Oxford Nanopore Technologies). An additional RAPID sequencing experiment was performed on a second *E. coli* gDNA sample to assess ONT sequencing variability. The second *E. coli* sample was sequenced on the MinION for 72 hrs, and data from the first 24 hr was used for comparisons.

To prepare ONT LIG libraries, gDNA was sheared to 20 kbp using g-TUBE (Covaris) and size-selected for fragments >10 kbp using the BluePippin system (Sage Science). Libraries were prepared with size-selected DNA using the Ligation Sequencing Kit SQK-LSK109 (Oxford Nanopore Technologies) according to the manufacturer’s protocol and including 1 μl DNA control sequence (DCS) in the master mix to validate library prep. Single 24 hr sequencing runs for *E. coli* and *D. ananassae* were performed with R9 MinION flowcells. An additional sequencing run was performed for *E. coli* using a FLO-MIN111 R10 MinION flowcell (Oxford Nanopore Technologies) that was produced without library size selection and shearing.

Base calling for all R9 runs was performed with Guppy v.3.1.5 using the ‘dna_r9.4.1_450bps_fast’ model. Base calling for the R10 run was performed with Guppy v.3.2.10 using the ‘dna_r10.3_450bps_fast’ model. DCS sequences were subsequently removed from ONT LIG fastq files using NanoLyse [49].

### PacBio libraries and sequencing

PacBio libraries were prepared using the SMRTbell Template Prep Kit 1.0/SMRTbell Express Template Prep Kit 2.0 (Pacific Biosciences). Genomic DNA was sheared to 20 kbp using g-TUBE, followed by DNA-damage repair and end-repair using polishing enzymes. Blunt-end ligation was used to create the SMRTbell template. Library fragments were size-selected using BluePippin. SMRTbell Polymerase Complex was created using DNA/Polymerase Binding Kit P6 v2 for RSII libraries and Sequel II Binding Kit 1.0 for Sequel II and HiFi libraries (Pacific Biosciences). PacBio RS II libraries were sequenced using DNA Sequencing Reagent Kit 4.0 v2 and RS II SMRT Cells v3 (Pacific Biosciences), with 4 hr movie length. Sequel II and HiFi libraries were sequenced using Sequel II Sequencing Plate 1.0 and SMRT Cells 8M (Pacific Biosciences), with 30 hr movie length.

### Illumina libraries and sequencing

llumina libraries for *E. coli* and *D. ananassae* were prepared using the KAPA HyperPrep kit (Kapa Biosystems, Wilmington, MA) using manufacturer’s instructions. Quantification of libraries was performed using the Quant-iT™ PicoGreen® dsDNA kit (Thermo Fisher Scientific). Library fragment size was assessed with the LabChip GX instrument (PerkinElmer). Paired end libraries (2×150 bp) were sequenced on an Illumina HiSeq4000 instrument (Illumina Inc.).

### *E. coli* genome assembly

Read length histograms were generated from the ‘readlength.sh’ script of bbtools v.38.47 [50] using 1 kbp bins. Reads from the ONT LIG and PacBio Sequel II libraries were randomly downsampled to approximate the sequencing depth of the PacBio RS II library using seqkit v.0.7.2 [51].

Assemblies were generated using Canu v.1.8 using default parameters and genomeSize=4.6m [52]. A custom script adapted from Chang & Larracuente [53] was used to iteratively polish genomes for five rounds using Pilon v.1.22 [54] with --minmq 10 and --fix bases. Separate polishing runs were performed with a) Illumina reads, b) corrected/trimmed long reads used in the Canu assembly, c) combined short and long reads, and d) PacBio HiFi reads. After polishing, circularization of the *E. coli* genome and candidate plasmids was attempted by the ‘minimus2’ command of Circlator v.1.5.5 [55] followed by *E. coli* genome rotation using the Circlator ‘fixstart’ command.

*E. coli* sequences were assembled separately using Unicycler v.0.4.8 [46], including long read assemblies and hybrid assemblies with Illumina data. The Unicycler pipeline includes a polishing step, performed here using Racon v.1.3.1 [56] for long read assemblies and Pilon using hybrid assemblies.

### Evaluation of *E. coli* genome assemblies

To detect plasmids in sequencing datasets, we submitted polished assemblies to the plasmid database PLSDB [27] using the mash dist search strategy with default parameters. To assess sequencing depth and estimated plasmid copy number, long reads were mapped to the consensus Unicycler genome using minimap2 v.2.1 [57] with default parameters and Illumina reads were mapped using bwa mem with -k 23 [58]. SAMtools v.1.9 [59] was used to filter out secondary and duplicate (Illumina only) alignments and calculate sequencing depth for each position. The estimated copy number for the *E. coli* genome and plasmids was determined by dividing the total number of bases mapping to each sequence by the total length of the sequence (**Table S2**). As a separate test of plasmid copy number, primers were designed for the *E. coli* genome, pMAR2, and p5217 from the Unicycler consensus assembly (**Table S16**) and pE2348-2 from NCBI (Genbank FM18070.1). Amplicons were quantified using the CFX384 Touch Real-Time Detection System and qPCR cycle threshold and melt curve values were obtained from CFX Maestro™ Software (Bio-Rad Laboratories Inc.). The mean cycle threshold (Ct) value for each sequence was calculated by averaging values from three replicates. ΔCt was calculated as the difference between the mean Ct value of the sequence of interest and the mean Ct genome. Estimated sequence copy number was calculated as 2^-ΔCt^ [60]. As a negative control, qPCR experiments also included samples with no template DNA.

The presence of highly conserved genes was determined using BUSCO v.3 [19] using the bacteria odb9 dataset from OrthoDB [61]. Following a strategy described in Wick *et al*. 2017 [20], assembly correctness was evaluated by generating assemblies of Illumina paired end data with ABySS v.2.1.1 [62] and separately with velvet v.1.2.10 [63]. Scripts from Wick *et al*. were used to identify assembled regions with >10 kbp length and 100% identity between the assembis as trusted contigs. Following this procedure, BLAST v.2.8.1 [64] was used to generate counts for mismatches, gap openings, and alignment lengths between trusted contigs and newly assembled *E. coli* genome sequences to determine overall correctness.

### Chimera detection

To evaluate chimeric read content in sequencing datasets, raw reads were aligned to the *E. coli* genome sequence using minimap2. Output files in paf format were used to identify putative chimeras using Alvis [22]. Using the -chimeras parameter, a read was called as chimeric when ≥90% of its length overlapped the consensus genome (-minChimeraCoveragePC 90) and two sub-alignments ≥10% of the total read length aligned to discordant regions of the genome (-minChimeraAlignmentPC 10). Since reads mapping to the two ends of the linear representation of the *E. coli* genome would be identified as chimeric, a second run of Alvis was performed with a rotated genome. Putative chimeras were calculated as the number of reads assigned as chimeras in both Alvis runs.

### *D. ananassae* genome assembly

The ONT RAPID, ONT LIG, and PacBio Sequel II libraries were assembled with Canu using genomeSize=240m and default assembly parameters. Hybrid assemblies were also generated with combined read data from Sequel II and one other library (ONT RAPID, ONT LIG, PacBio RS II). The Sequel II + ONT LIG assembly was iteratively polished for five rounds using Pilon using Illumina reads, corrected long reads, and PacBio HiFi reads.

### Dana.UMIGS genome assembly

A new *D. ananassae* reference assembly (referred to as Dana.UMIGS) was generated separately to enable comparisons between test assemblies produced in this study. The PacBio and ONT libraries were assembled with Canu using genomeSize=240m, corOutCoverage=80, and other default assembly parameters. The output assembly was polished for two rounds with Arrow v.2.3.3 (SMRTTools v7) using Sequel II read data. The Sequel II reads were also assembled with Flye v. 2.7.1 [65] in raw pacbio mode using -g 240m and --asm-coverage 60 and included a polishing step. Following the merger of the Canu (query) and Flye (reference) assemblies with quickmerge v.0.3 [66] using conservative parameters (-ml 5000000 −l 20000), the resulting merging events were manually inspected. Unsupported merge events were split into constitutive contigs. The final merged assembly was polished for one round with Sequel II reads using Arrow and five rounds with HiFi reads using Pilon. To assess the presence of duplicated content, PacBio HiFi reads were mapped to the assembly using minimap2 and a histogram of sequencing depth across the genome was produced using purge_haplotigs v.1.1.1 [67]. After manually inspecting assembly contigs classified as ‘junk’ or ‘suspect’ using a low depth cutoff of 10 and a high depth cutoff of 200, we removed 86 contigs corresponding to bacterial contaminants, assembly artefacts, and erroneous duplications in the assembly.

### Anchoring Dana.UMIGS contigs

To identify contigs in the Dana.UMIGS assembly corresponding to the major euchromatic chromosomes (X, 2, 3), regions with known positions on chromosome arms were extracted from the caf1 assembly. Coordinates of the caf1 regions were reported previously [31]. Dana.UMIGS assembly contigs were searched for caf1 sequences using BLASTN. After initial searches using default parameters indicated the presence of high-quality matches as the first hits for each caf1 query, a second BLASTN search was conducted to retain the single best hit for each query (-max_target_seqs 1 -max_hsps 1). The positions of loci in the Dana.UMIGS assembly were plotted using a custom script in R.

To anchor contigs to chromosome Y, two male and two female 2×x150 bp Illumina libraries were prepared and sequenced using the same methods as the mixed sex *D. ananassae* Illumina library. Reads were randomly downsampled to ~150X depth using seqkit and mapped to the Dana.UMIGS assembly using bwa mem. Duplicate reads were removed with Picard MarkDuplicates (Broad Institute) and the sequencing depth for each position in the genome was determined with SAMtools depth while removing low-quality mappings (-Q 10). A script adapted from Chang & Larracuente [53] was used to split the genome into 10 kbp windows and determine the median of the female/male sequencing depth ratio for each window. Contigs were assigned as putative Y contigs as having (a) at least one window with a median female/male ratio of zero, (b) ≥80% of its windows with median female/male ratios below 0.05, and (c) conditions (a) and (b) met for two female-male replicates.

To anchor contigs to chromosome 4, Dana.UMIGS contigs were aligned to caf1 assembly contigs previously assigned to chromosome 4 [34] using nucmer with -l 1000 and --maxmatch. Chromosome 4 contigs containing LGT from the fly’s Wolbachia endosymbiont (*w*Ana) were anchored by aligning Dana.UMIGS contigs to the previously assembled *w*Ana genome [68] with -l 1000 and --maxmatch.

### Evaluation of *D. ananassae* genome assemblies

To generate assembly statistics, assembly contigs were evaluated using QUAST-LG [69] with the Dana.UMIGS assembly. NGX plots were generated with a script adapted from the Assemblathon 2 paper with 231 Mbp as an estimated genome size [70]. BUSCO searches were conducted using the metazoa odb9 dataset. For comparisons to published *D. ananassae* assemblies, the same analyses were performed on the caf1 genome assembly (Genbank GCA_000005115.1) published as part of the *Drosophila* 12 Genomes Project [16] and a genome assembled by Miller *et al*. [17] using ONT sequencing.

To evaluate contiguity of the euchromatic chromosome arms in assemblies produced from single long read libraries, contigs corresponding to chromosome X, 2, and 3 were extracted from each assembly using BLASTN-based searches as described above for Dana.UMIGS. Dana.UMIGS contigs were aligned to contig sets with nucmer using -l 200 and - -maxmatch.

To evaluate contiguity of heterochromatic regions, the contig sets that were not anchored to euchromatic chromosomes were extracted from single library assemblies. Summary statistics for heterochromatic contigs were produced using QUAST-LG without specifying a reference genome.

### DNA modification

Detection of DNA methylation using *E. coli* PacBio libraries was assessed with PacBio SMRT Tools. Differences in RS II and Sequel II libraries necessitated the use of similar pipelines (*i.e.,* PacBio base modification pipeline) on different software releases (RS II: SMRT Link v.7; Sequel II: SMRT Link v.8). Reads were mapped to the *E. coli* genome with pbmm2 and detection of DNA methylation signatures was performed with ipdSummary using --identify m4C,m6A,m5C_TET to search for m4C, m6A, and m5C modifications, respectively. Highly modified motifs were identified with motifMaker. The distribution of modification QV scores for the four nucleotide bases was produced by the SMRT Tools pipeline and an appropriate modification QV cutoff was determined.

Detection of DNA methylation using *E. coli* ONT libraries was assessed with Tombo v.1.5 [26] using the *de novo* model for modified base detection. The dampened fraction of reads supporting each modification event was produced with tombo text_output. The presence of DNA modification at specific motifs was assessed using the tombo plot motif_with_stats command by plotting the dampened fraction values for up to 10,000 genomic sites containing the motif of interest. ROC curves for detected GATC and CCWGG motifs in ONT libraries were produced with the tombo plot roc command.

DNA methylation detection in the *D. ananassae* ONT LIG library was conducted using Tombo similar to the methods described in *E. coli*. Given the lack of known methylated motifs in *D. ananassae*, *de novo* modified base detection was followed by the extraction of 1000 regions showing the largest estimated dampened fraction of modified bases using the tombo text_output command. The presence of overrepresented motifs in candidate modified regions was evaluated with MEME v.4.12.0 using the parameters -dna -mod zoops -nmotifs 50.

## AVAILABILITY OF DATA AND MATERIALS

All the data supporting the conclusions of this article have been deposited in Genbank/EMBL/DDBJ Sequence Read Archive under BioProject PRJNA602597. Additional figures and tables supporting the conclusion of the article are available as Supplementary Information. All commands and scripts used in the study are available at https://github.com/Dunning-Hotopp-Lab/Ecoli-Dana-LongReads.

## ACKNOWLEDGEMENTS

Not applicable.

## FUNDING

This project has been funded by the National Institute of Allergy and Infectious Diseases, National Institutes of Health, Department of Health and Human Services under grant U19AI110820. JCDH and EST are also supported by an NIH Director’s Transformative Research Award (R01CA206188).

## ETHICS DECLARATIONS

### Ethics approval and consent to participate

Not applicable.

### Consent for publication

Not applicable.

### Competing interests

We have no competing financial interests.

### Contributions

EST and JCDH wrote the manuscript. All authors read and edited the manuscript. JM and DAR prepared and provided *E. coli* DNA samples, MG and JCDH prepared and provided *D. ananassae* samples. MG and BCS prepared sequencing libraries. BCS conducted PCR and qPCR validation experiments. MG, BS, XZ, LJT, LS, and JCDH sequenced or oversaw sequencing. EST performed bioinformatic analysis and generated figures. RB obtained and modified publication-quality polytene map images and prepared sequencing data submission to SRA. JCDH, DAR, LJT and LS designed and supervised the study.

## SUPPLEMENTARY FIGURE LEGENDS

**Figure S1. Read composition of subsampled *E. coli* long read libraries**.

The ONT LIG and PacBio Sequel II reads were randomly sampled to approximate the sequencing depth of the PacBio RS II library using seqkit [51]. Single end reads were placed into 1 kbp read length bins. Read counts and sequenced bases were calculated for each bin using the readlength.sh script of bbtools [50]. **A.** Absolute values for read counts and sequenced bases are plotted for each read length bin. **B.** Read counts and sequenced bases expressed as percentages of the total for each library are plotted for each read length bin. Vertical dotted lines correspond to maximum read length for each library.

**Figure S2. Dot plot alignment of the *E. coli de novo* Unicycler assemblies**.

For each library, hybrid assemblies were generated using Unicycler with short Illumina reads alongside long reads and subsequently polished with short reads. The assemblies were concatenated into a single file and a self-self dot plot was constructed using NUCmer with the maxmatch parameter and visualized using mummerplot. The colors of alignments represent percent similarity between sequences.

**Figure S3. Validation of deletion region in *E. coli* E2348/69**.

**A.** Hybrid assemblies using Illumina sequencing with ONT RAPID, ONT LIG, PacBio RS II, or PacBio Sequel II were aligned to a previously reported assembly for this strain (Genbank GCA_000026545.1) [15] using nucmer with the - -maxmatch parameter. Colors of alignments indicate the percent similarity of alignments. The grey box outlines a ~16 kbp region that is present in the published sequence of *E. coli* but is missing in the genomes assembled in this study. **B.** Agarose gel electrophoresis of gene sequences in *E. coli* E2348/69. Primers were designed for the following sequences: NADP: NADP-dependent phosphogluconate dehydrogenase (WP_000043439.1) FLP: M-antigen undecaprenyl disphosphate flippase (WP_000058470.1), WcaM: colanic acid biosynthesis protein WcaM (WP_001115987.1), Wcz: tyrosine-protein kinase Wzc (WP_000137154.1), 18S(−): *D. ananassae* 18S rRNA primers with no DNA, 18S(+): *D. ananassae* 18S rRNA primers with fly DNA. M: Marker. The putative ~16 kbp deletion region includes FLP and WcaM while the NADP and Wcz genes on opposite sides of this region are present in *E. coli* assemblies generated in this study.

**Figure S4. Visualization of chimeric reads mapped to the *E. coli* genome**.

Reads with blue outlines are supplementary alignments. Blue, red, green, and orange ticks represent SNPs relative to the reference genome, black dots represent indels. From top to bottom: ONT RAPID, ONT LIG, PacBio RS II, PacBio Sequel II. Split reads mapping to distinctive regions were present (connected by red lines in the figure), which represent chimeric reads. There was no apparent evidence of supplementary alignments immediately adjacent to primary alignments representing erroneous chimera assignment. Screenshot captured in IGV [74].

**Figure S5. Distribution of modification QV values in *E. coli* PacBio libraries**.

PacBio Sequel II and RS II reads were processed by the SMRT Tools DNA modification pipeline, including read mapping and DNA modification detection using interpulse duration (IPD) values. Each methylated site is assigned a modification quality value (QV) score based on differences between observed and expected IPD values. Distributions of modification QV values are plotted for each of the four nucleotide bases.

**Figure S6. Assessment of m6A modification in *E. coli* E2348/69 DNA motifs**.

DNA methylation at YTCAN^6^GTNG, and CYYAN^7^RTGA, and ATGCAT motifs are assessed using ONT LIG sequencing. Top: An example motif is shown, with red lines displaying individual reads mapped to the region. The expected raw signal distribution using a canonical base model (=unmethylated DNA) is shown in grey. The location of known methylation in *E. coli* is highlighted in yellow. Bottom: The fraction of reads supporting a modification event is reported for each position in the motif, and the distribution of proportions are shown. Higher values indicate the motif is more ubiquitously methylated in the *E. coli* genome. Distributions are shown for 242 YTCAN^6^GTNG motifs, 248 CYYAN^7^RTGA motifs, and 66 ATGCAT motifs.

**Figure S7. qPCR amplification of *E. coli* sequences**.

**A.** qPCR amplification of *E. coli* sequences. The horizontal black line represents the threshold set by Bio-Rad CFX384 for calling Ct values (RFU = 4000). Vertical dotted lines represent mean Ct values for each target sequence. **B.** Melt curves from qPCR of *E. coli* genome and plasmids. NTC – negative template control.

**Figure S8. Read composition of *D. ananassae* long read libraries**.

Reads were binned in 1 kbp intervals, and various statistics were calculated using readlength.sh script of the bbtools package [50]. Top: Read counts are plotted for each read length bin.

Bottom: Cumulative percentage of bases sequenced is plotted for each read length bin. Vertical dotted lines correspond to maximum read length for each library. See Table S1 for summary of read libraries.

**Figure S9. NGX plot of *D. ananassae* genome assemblies**.

Plot of NGX values for *D. ananassae* assemblies produced in this study. Each NGX value represents the shortest contig length when summed with all larger contigs totaling X% of the estimated genome size. Assemblies produced here were compared to two previous assemblies of *D. ananassae* [16, 17], and the 231 Mbp assembly size (including gaps) of the Drosophila 12 Genomes Consortium caf1 assembly was used to calculate NGX values.

**Data S1. Methylated motifs in D. ananassae**

ONT LIG reads were evaluated for evidence of DNA methylation using Tombo with the ONT LIG + Sequel II assembly as a reference. A total of 1,000 regions showing the strongest evidence for methylation were extracted using tombo text_output and screened for enriched motifs using MEME.

## Notes

### Competing Interest Statement

The authors have declared no competing interest.

