## Supplementary figures and images for "Comparison of long read sequencing technologies in resolving bacteria and fly genomes"

### Figure S1

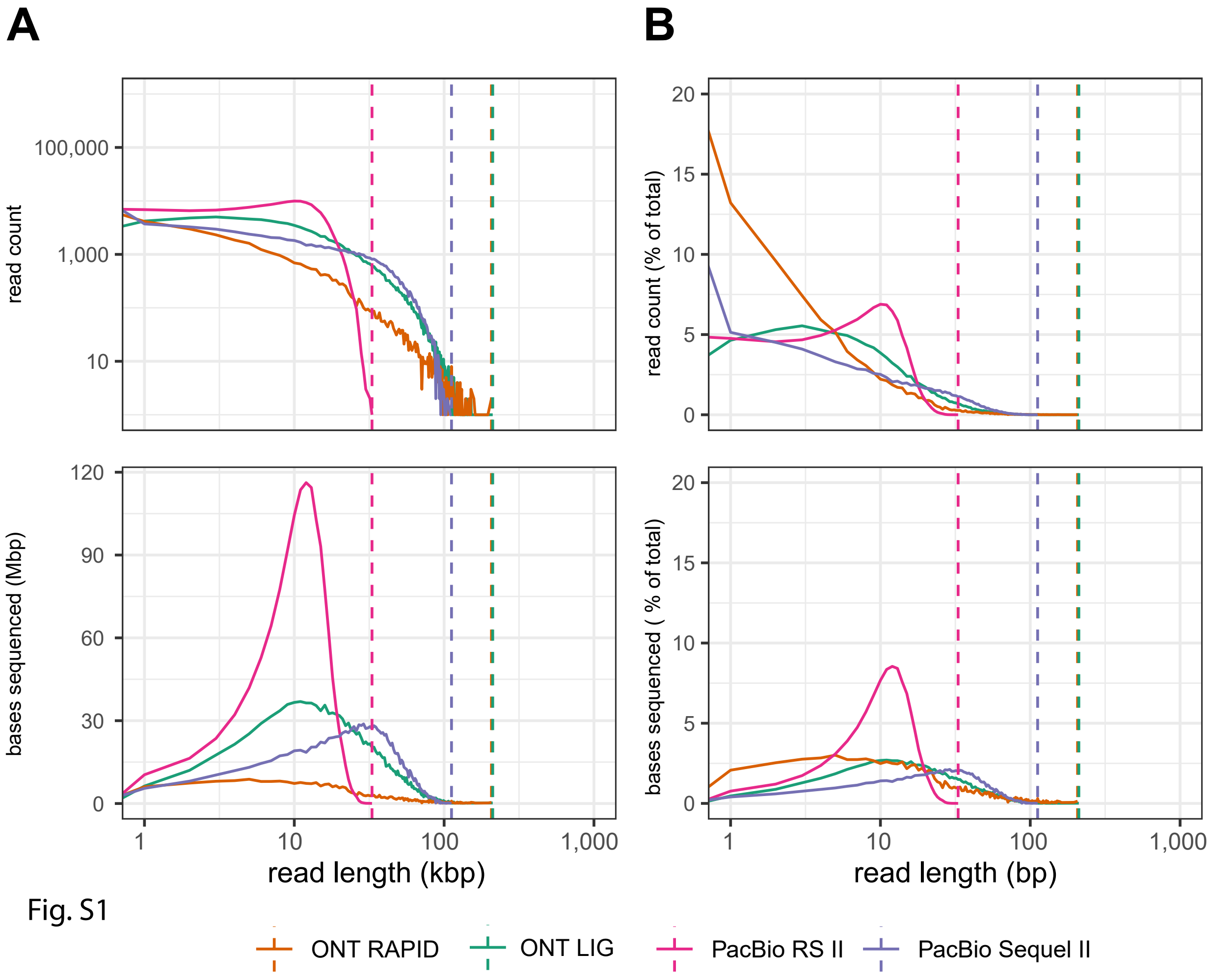

### Figure S2

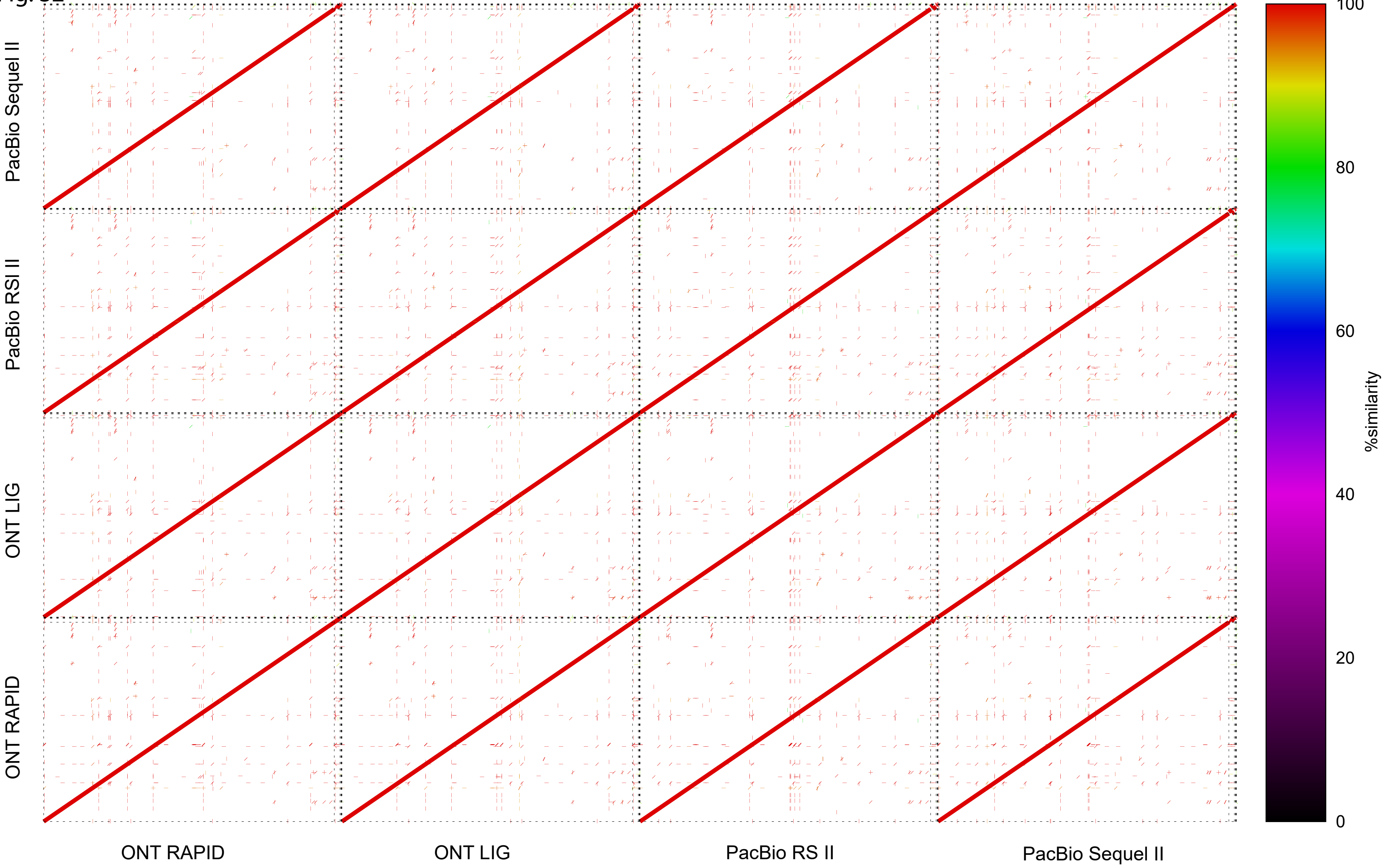

### Figure S3

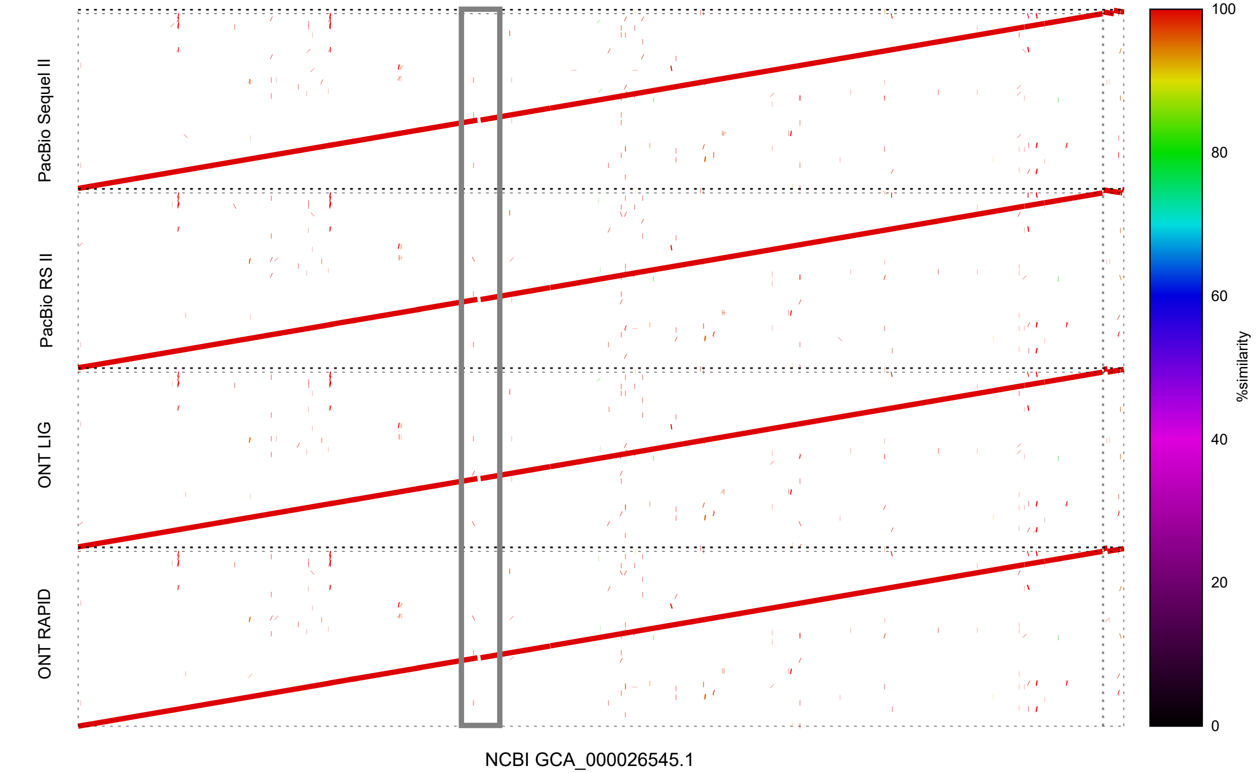

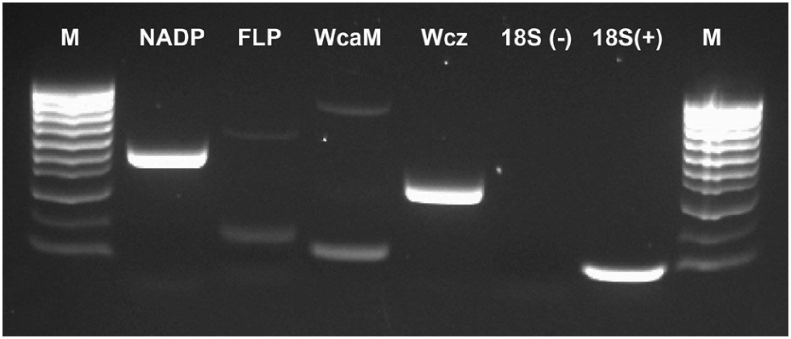

### Figure S4

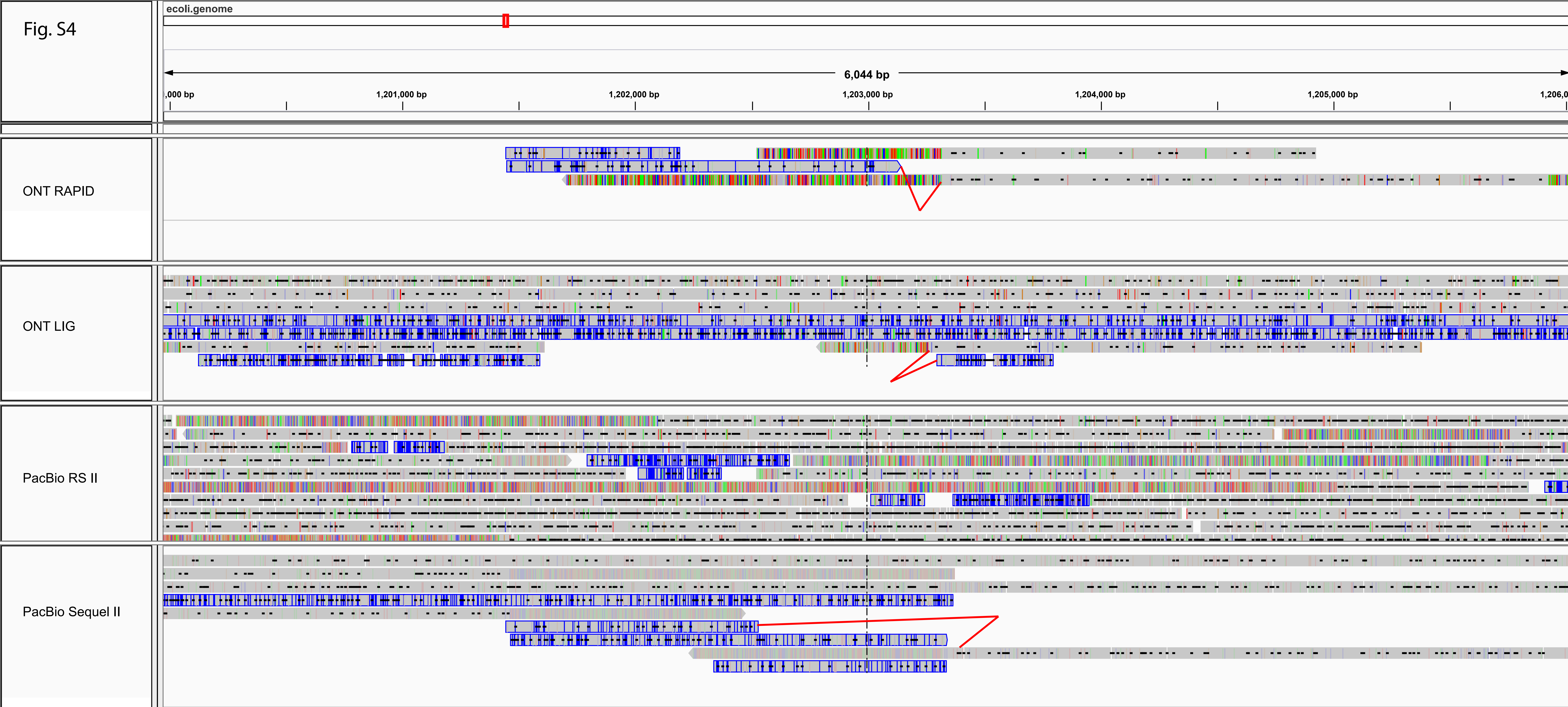

### Figure S5

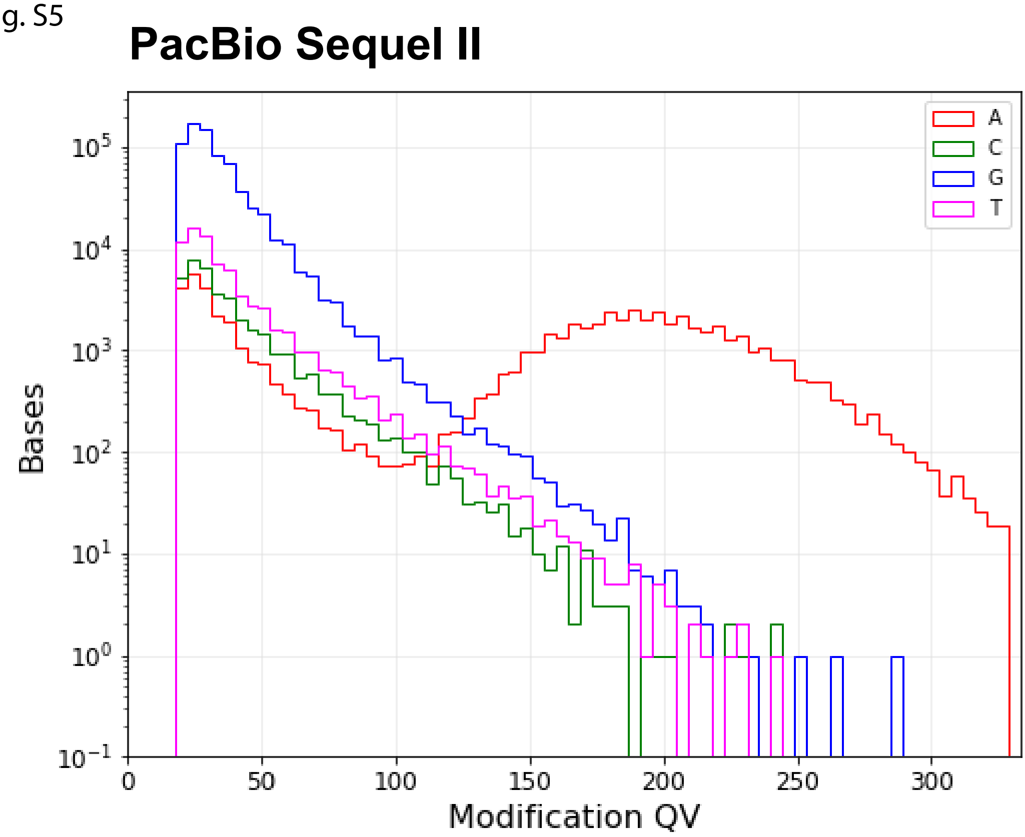

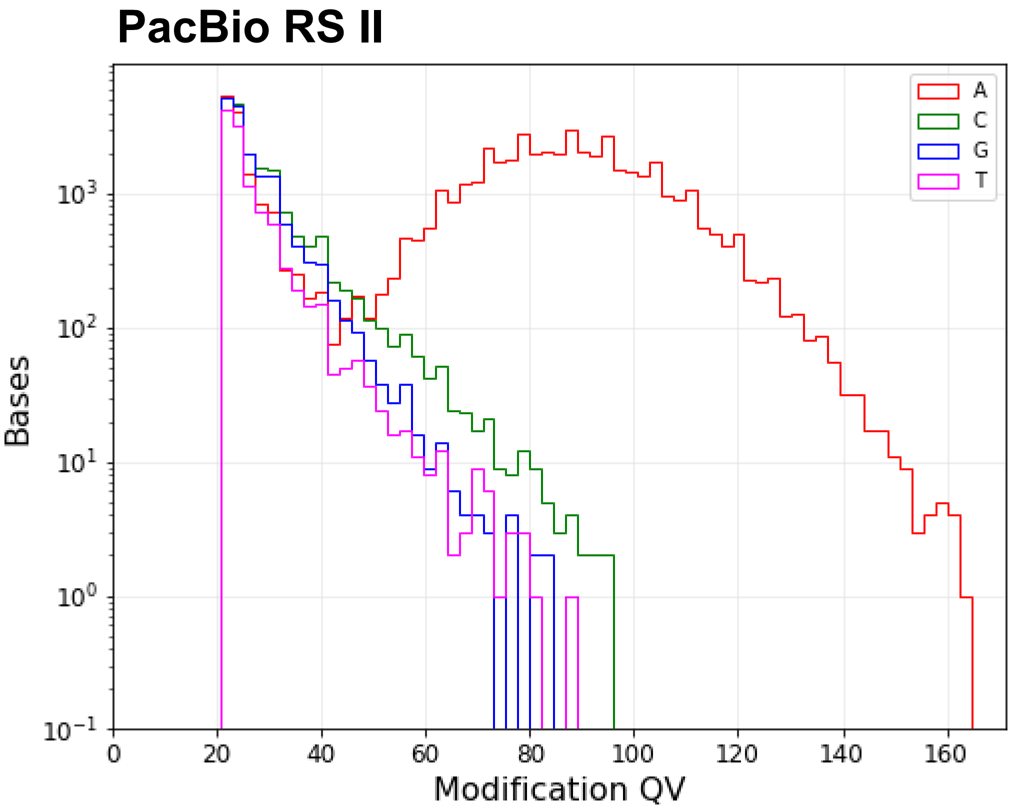

### Figure S6

Fig. S6

#### YTCAN$^6$GTNG

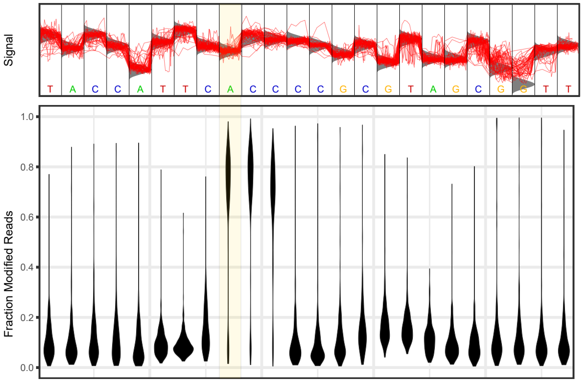


#### CYYAN$^7$RTGA

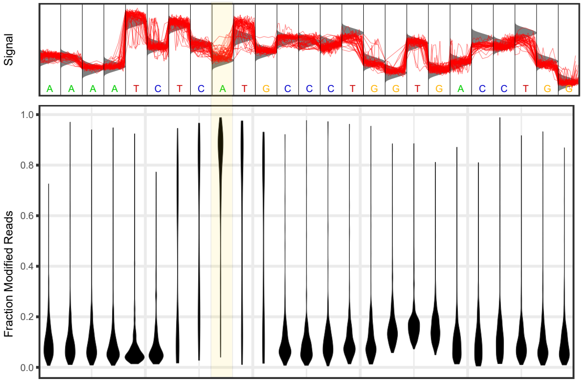


#### ATGCAT

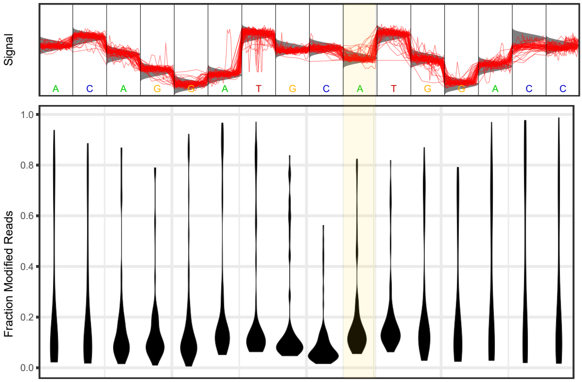

### Figure S7

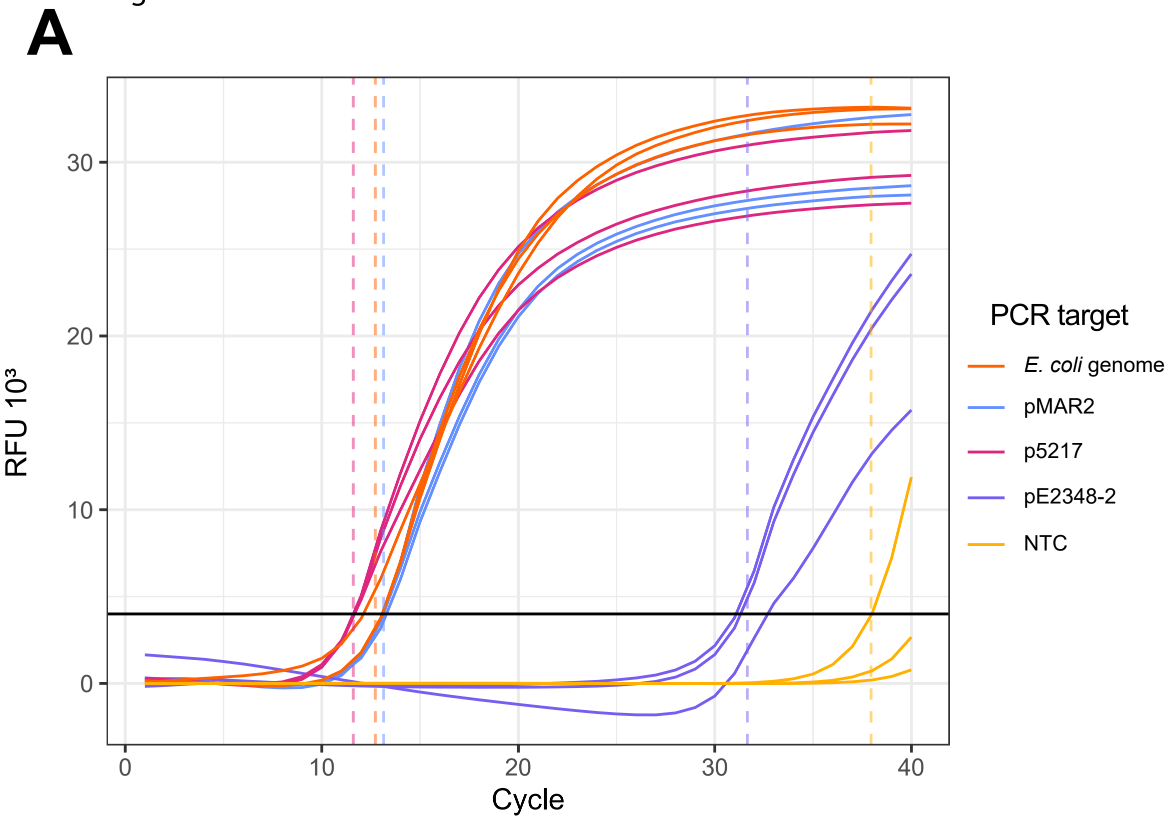

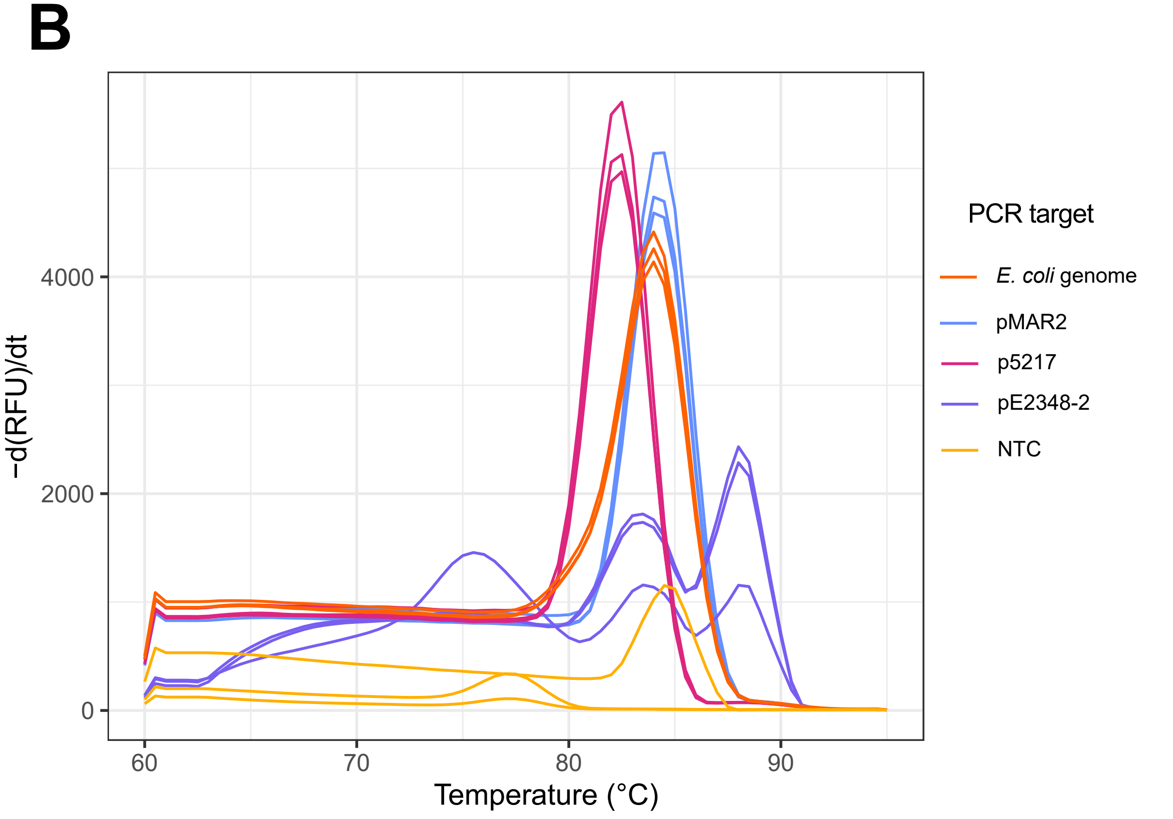

### Figure S8

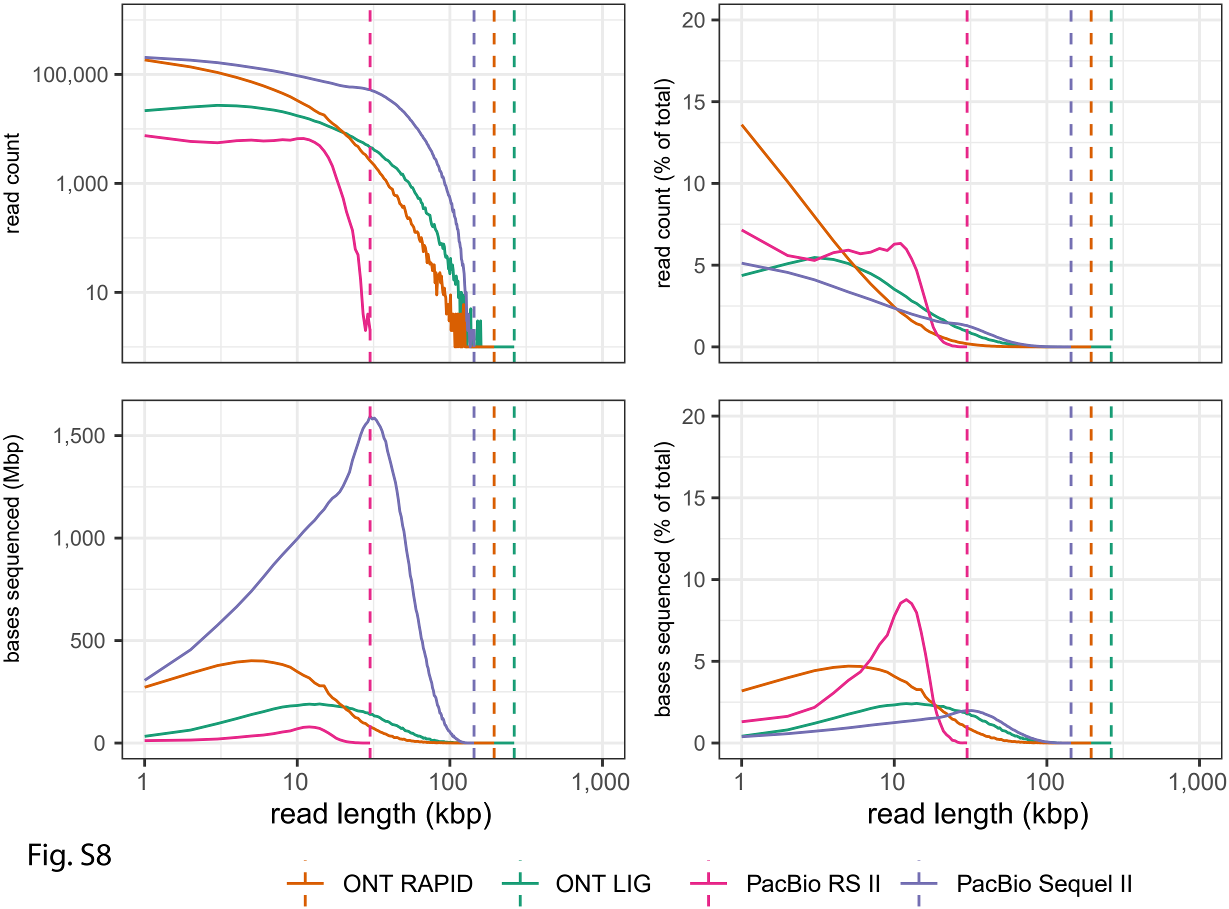

### Figure S9

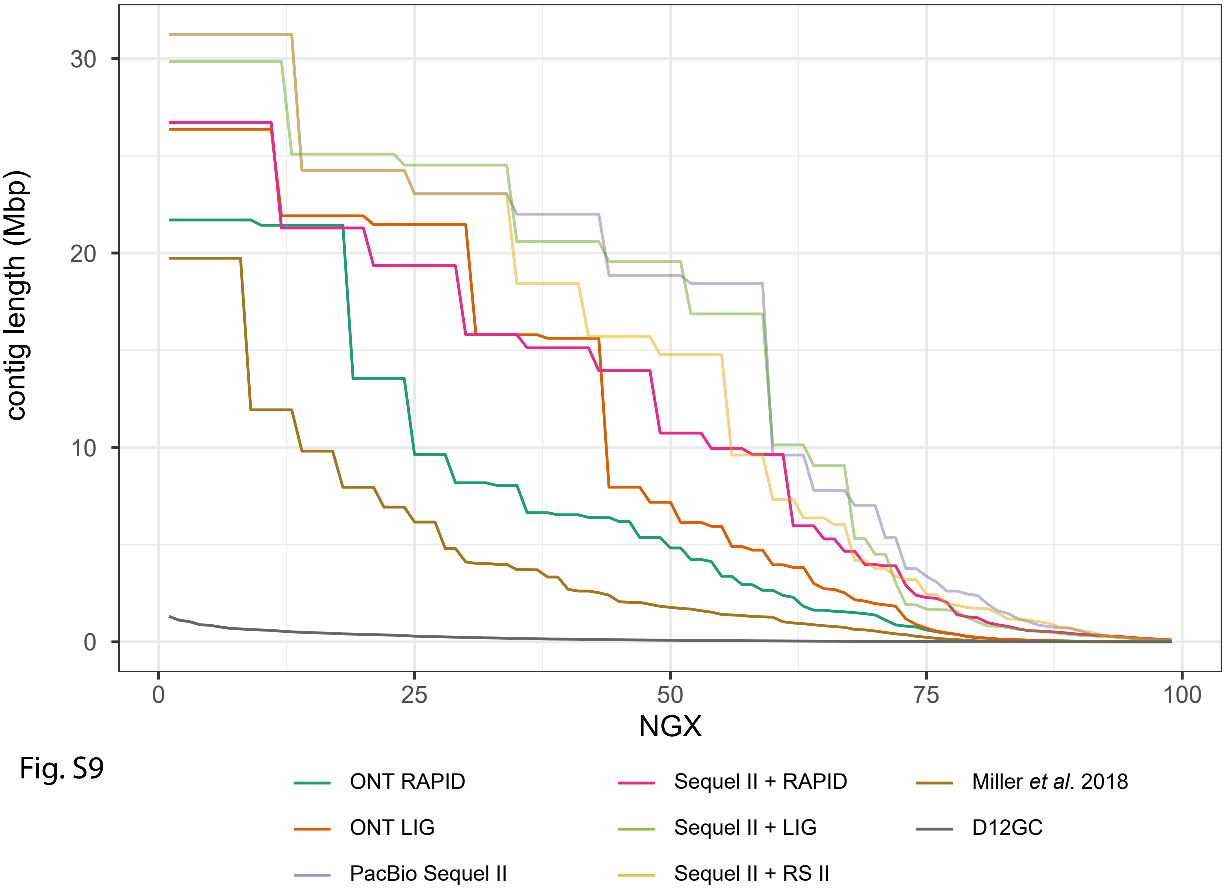
